# Factors Influencing Phenomic Prediction: A Case Study on a Large Sorghum BCNAM Population

**DOI:** 10.1101/2025.02.04.636400

**Authors:** Clément Bienvenu, Vincent Garin, Nicolas Salas, Korotimi Théra, Mohamed Lamine Tekete, Madathiparambil Chandran Sarathjith, Chiaka Diallo, Angélique Berger, Caroline Calatayud, Fabien De Bellis, Jean-François Rami, Michel Vaksmann, Vincent Segura, David Pot, Hugues de Verdal

## Abstract

Plant breeding efficiency is crucial to develop varieties able to cope with climate change and support food and feed value chains. Genomic prediction (GP) has been a major step in increasing this efficiency and is now routinely used in breeding programs. Recently, phenomic prediction (PP) has gained attention as a promising complementary approach to GP, further increasing the breeding programs’ efficiency. Factors impacting the predictive ability (PA) of PP have been studied on many species but are not fully clarified. In this context, we studied the impacts of spectra pre-processing, prediction methods, population structure, training set size, NIRS acquisition environment and wavelength selection on a large multi-parental sorghum population including 2498 genotypes.

Our results show that PP can compete with GP, that it is less affected by population structure, and can reach its maximal PA with smaller training sets than GP, but its performances are trait dependant. We also show that NIRS can be acquired in a reference environment to perform prediction in other environments and that it is possible to randomly select as little as 10 wavelengths to perform predictions. Finally, we show that spectra pre-processing, and statistical methods have a limited and unclear impact on PA. Our study confirms that PP is a relevant trait prediction method that deserves attention to optimize breeding schemes. The main challenges for the future will be to better understand the information contained in the spectra and disentangle their genetic and proxy components to optimize the use of PP in breeding programs.

**Key message:** Phenomic prediction is promising for sorghum breeding. Geneticists’ methods may not be suited to optimally extract spectral information.

## Introduction

Plant breeding plays a key role in food and feed production, by allowing the development of varieties adapted to the needs and stakes of the value chain stakeholders. Thus, improving breeding programs’ efficiency is crucial to guarantee food security especially in the context of climate change (Tester and Langridge, 2010). By referring to the breeder’s equation (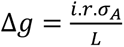 (Lynch et al., 1998)), a breeding program can be improved by increasing selection intensity (*i*), better estimating breeding values (*r*), screening more diversity (σ*_A_*), or reducing the time and cost of field experiments (*L*).

Genomic prediction (GP) has been conceptualized in the late 1990s and is a method to estimate breeding values using only molecular markers, thus reducing the need for expensive field experiments once a predictive model is set (Meuwissen et al., 2001). It has been routinely implemented in a wide range of species to better estimate breeding values and increase selection intensities. Through its two decades of existence, GP has been tested in a variety of contexts, and some key factors impacting its reliability have been studied: genotyping density, genetic architecture of the predicted trait, training set size, relatedness between training and validation sets, statistical methods and modelling, and the importance of genotype-by-environment (GxE) interactions (Lado et al., 2016; Lorenz et al., 2011).

Recently, (Rincent et al., 2018) suggested an innovative approach called phenomic prediction (PP) to further reduce the costs of breeding values estimations. PP globally mimics GP but uses near infrared reflectance spectroscopy (NIRS) of plant tissues or canopies as predictors instead of molecular markers. It is low cost, high throughput, non-destructive and often ready to be implemented in breeding programs that already gather infrared spectroscopy data for other uses. Considering a broad definition of PP, the technique has been implemented in mainly three forms: (i) with tissue or plant organ based spectra using a wide range of wavelengths, (ii) with canopy based spectral measurements often done with unmanned aerial vehicles (UAVs) carrying multi- or hyper-spectral cameras covering a narrower range of wavelengths in the visible and near infrared, (iii) with vegetation indices (VI) also based on canopy measurement but with spectra pre-processing to calculate VIs that play the role of the predictor. Phenomic predictions have been reported on a wide range of plant species, including annual crops such as wheat (Dallinger et al., 2023), maize (Adak et al., 2022), soybean (Zhu et al., 2021), rye (Galán et al., 2020), triticale (Zhu et al., 2022), mungbean (Fumia et al., 2023b), rapeseed (Roscher-Ehrig et al., 2024), potato (Maggiorelli et al., 2024), pepper (Fumia et al., 2023a), and rice (Verdal et al., 2024), as well as perennials such as alfalfa (Feng et al., 2020), sugar cane (Gonçalves et al., 2020), coffee (Adunola et al., 2024), grapevine (Brault et al., 2022), and even forest trees such as poplar (Rincent et al., 2018), slash pine (Li et al., 2023), and eucalyptus (Mora-Poblete et al., 2024). Phenomic predictions have also recently been reported for dairy sheep (Machefert et al., 2024).

Key factors impacting PP reliability are the same as GP and some of those factors have also been studied for PP but less extensively. Moreover, some factors specific to PP also deserved to be studied, namely, spectra pre-processing and number of wavelengths used. In addition, contrarily to genotyping data, which are fixed for a given genotype, NIRS data captures environmental information. Thus, environment of NIRS acquisition (phenological stage, tissue, and environmental context…), and environmental relatedness between training and test sets are also impacting the reliability of PP. As spectra contain genetic and environmental information, it should be advantageous to replicate measurements and better isolate the genetic part while also taking advantage of GxE information.

Overall, it has been found that PP can perform similarly to GP in many different species (Adunola et al., 2024; Brault et al., 2022; Dallinger et al., 2023; Rincent et al., 2018; Robert et al., 2022b, 2022a; Roscher-Ehrig et al., 2024; Thapa et al., 2024; Weiß et al., 2022; Zhu et al., 2022, 2021) and that statistical methods do not seem to impact PP predictive abilities (PA) (Roscher-Ehrig et al., 2024; Zhu et al., 2021), but non-linear methods and deep learning algorithms may provide some improvement over more classical linear models (Cuevas et al., 2019; Mora-Poblete et al., 2024). PP is also less sensitive than GP to population structure (Laurençon et al., 2024; Roscher-Ehrig et al., 2024; Weiß et al., 2022; Zhu et al., 2022, 2021), and it requires fewer genotypes in the training population to reach its maximal PA (Dallinger et al., 2023; Zhu et al., 2022, 2021). The quantitative nature of the trait (high/low number of QTL) seems to affect PP PA, with better performance for complex traits such as yield compared to GP (Roscher-Ehrig et al., 2024; Zhu et al., 2022). It also seems that selecting wavelength based on LASSO weights or PLSR loadings can greatly reduce the number of wavelengths needed with little impact on PA (DeSalvio et al., 2024; Zhu et al., 2021). One hypothesis to explain these performances of PP is that each wavelength would capture the effects of many genes at the same time (Zhu et al., 2021). Thus, a single wavelength would be more informative than a single SNP, which is likely due to their continuous nature. From this statement, it seems possible that PP can perform as high as GP with only a few dozen of wavelengths as they already cover a large part of the genome (Zhu et al., 2021). It is also coherent with the fact that PP can outperform GP on complex traits but not on mono-/oligo-genic traits. Indeed, by capturing the effects of many genes at the same time, wavelengths can also capture non additive information such as epistasis or GxE interaction. As mono-/oligo-genic traits are driven by a few QTL with strong additive effects explaining a large part of the phenotypic variance, PP could differentiate lines differing at a few loci only if they directly affect a wavelength of the spectrum. On the contrary, traits less dependent on addictive effects could be better predicted by PP (Zhu et al., 2022). Finally, this hypothesis can also explain the need for smaller training sets, as more information is carried by each spectrum, less spectra are required to train a model (Zhu et al., 2022).

More specifically to PP, NIRS environment acquisition has an impact on PA, with a drop of performance when spectra of the training and validation sets are not from the same environment (Zhu et al., 2021) likely due to the environmental information in the spectra. Combining spectra from different environments, phenological stages, or tissue may improve prediction abilities (Rincent et al., 2018; Robert et al., 2022a, 2022b) maybe because different tissues/time of acquisition have different gene expression profiles and can complement each other’s missing genetic information. It has also been shown that having spectra in a reference environment to predict genotypes’ performances in other environments is feasible (Brault et al., 2022; Rincent et al., 2018; Roscher-Ehrig et al., 2024; Zhu et al., 2022, 2021) which supports the hypothesis that NIRS can capture genetic information and relatedness between genotypes. Lastly, PP has been tested in many scenarios relevant for breeding programs with best results for sparse testing, poorest results for predicting unknown genotypes in unknown environments and intermediate and contrasted results across studies for unknown genotypes in known environments and known genotypes in unknown environments (Adak et al., 2022; Adunola et al., 2024; DeSalvio et al., 2024; Lane et al., 2020; Robert et al., 2022b).

Sorghum (*Sorghum bicolor*) is the fifth cereal in the world after corn, wheat, rice and barley with a total production of 57 MT produced in 2022. Africa is the biggest producer accounting for 51% of the world production (“FAOSTAT,” 2024). It is a very versatile crop as its use include food (raw and processed), cattle feed (grain and fodder), and energy production (as a bio-ethanol source, mostly in the USA) and even phytoremediation of cadmium-contaminated soils (Liu et al., 2020). Moreover, its tolerance to drought, low input levels or salinity makes it a very interesting crop to cope with climate change (Hossain et al., 2022). Despite being an important crop, PP has never been tested on this species which could beneficiate a lot from low-cost breeding tools as it is mainly used in developing countries.

The present study was conducted on a large (multi parental) sorghum BCNAM population (Garin et al., 2024) from which a subset of two recurrent parents and 22 donor parents grown during two years in 4 environments was considered. The strong genetic structure of the population, and the several environments in which it was phenotyped make it particularly suited to study the impact of population structure and NIRS environment acquisition on PP PA. Moreover, multi parental populations are relevant for breeding programs but were not frequently used in PP studies. In this context, we investigated the potential of PP for sorghum breeding by studying the factors impacting PP: training set size, population structure, NIRS acquisition environment, wavelength selection, spectra pre-processing, and statistical method.

## Materials and methods

The plant material, phenotypic and genomic data acquisition were described in Garin et al., (2024). We briefly recall those points in the following sections. Thus, the “Plant material”, “Phenotypic data” and “Genotypic data” sections are taken from Garin et al., (2024) with adaptations to fit the analysis made in this study. The general framework of the study is schematized in **Fig. 1**.

**Fig. 1.**
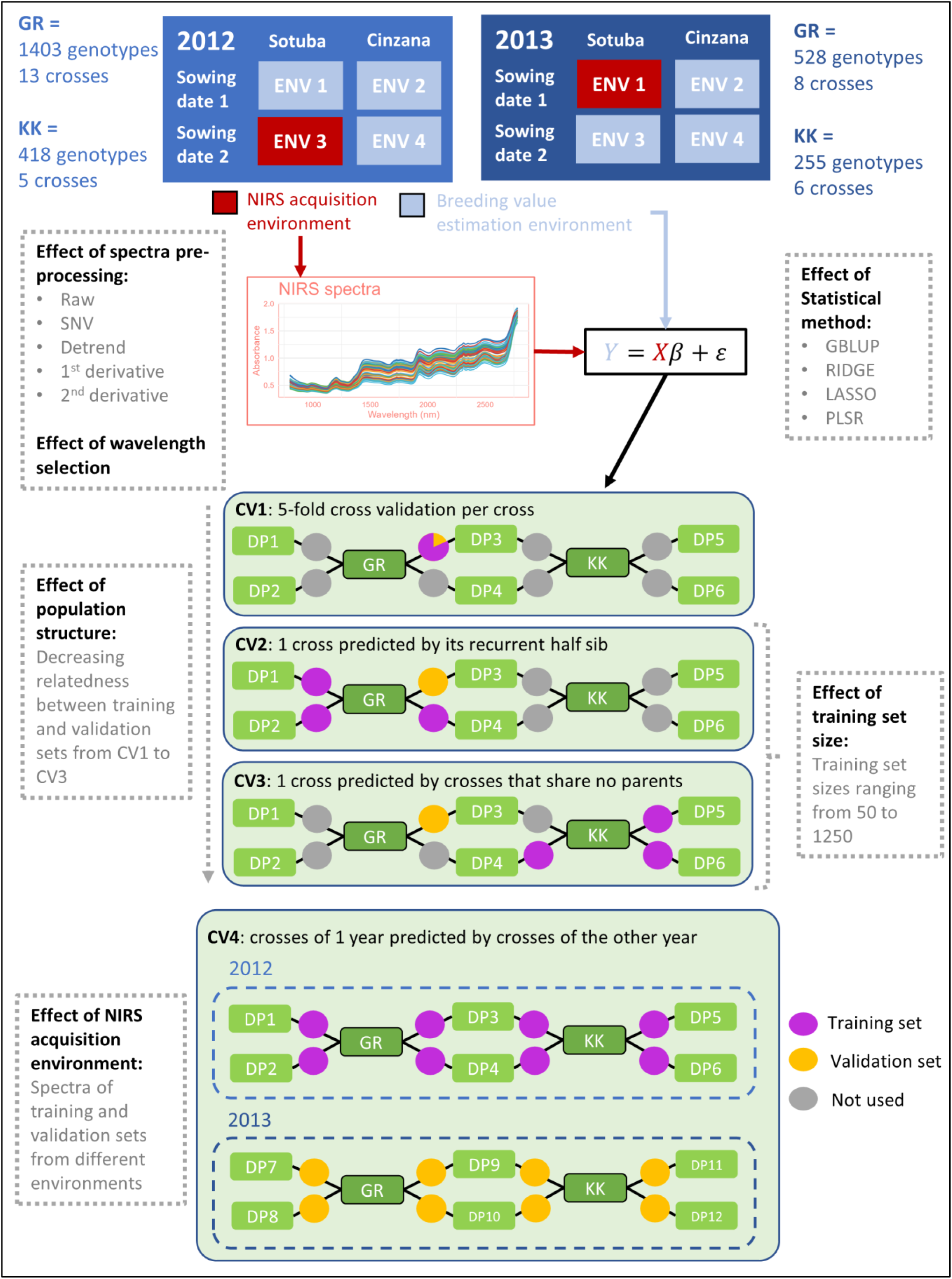
Schematic representation of the data, the cross-validation scenarios, and the factors studied. GR = Grinkan, KK = Kenin Keni, DP = donor parent

### Plant material

The WCA-BCNAM population (West and Central Africa Back Cross Nested Association Mapping) is composed of three populations obtained by crossing three elite recurrent parents to 24 donor parents, backcrossing their F1 hybrids to obtain BC1 families which were then selfed for three generations to obtain BC1F4 genotypes. Data were acquired on BC1F3:4 generations. NIRS data was acquired in only two of these populations which were retained in this study: Grinkan (GR) and Kenin-Keni (KK). The recurrent parents are elite lines selected in Mali through farmer variety testing. GR was developed through pedigree breeding methods. KK was derived from a recurrent selection population involving local parents of different botanical types (Leroy et al., 2014). These two recurrent parents were chosen for their productivity, their adaptation to soil and climate, and their resistance to major biotic and abiotic stresses, but they have poor grain quality and mold susceptibility (GR), and low yield and yield stability (KK). The 24 donor parents cover diverse racial (Guinea, Caudatum, Durra (Harlan and de Wet, 1972)) and geographical origins. They are characterized by key adaptive traits like height, maturity, and photoperiod sensitivity. Those parents were also selected for traits like tolerance to S*triga hermonthica*, soil phosphorus deficiency and/or drought, and good grain quality that could increase farmer acceptance (see Garin et al., 2024, Table 1).

**Table 1.** Summary of scenarios and training set sizes used to study each factor.

| Analysis | CV scenario | 2012 training set size | 2013 training set size |
| --- | --- | --- | --- |
| PP vs GP | CV2 | 1000 | 450 |
| Pre-treatment |  |  |  |
| Statistical method |  |  |  |
| Wavelength selection |  |  |  |
| Training set size | CV2, CV3 | 50, 100, 150, ... , 1250 | 50, 100, 150, ... , 400 |
| Population structure | CV1, CV2, CV3 | 100 | ∅ |
| NIRS acquisition environment | CV4 | 600 | 600 |

### Phenotypic data

Genotypes were phenotyped during two years (2012 and 2013) in Mali at two sites: Sotuba (12.65, −7.93) and Cinzana (13.25, −5.96) with two sowing dates (Sowing 1: end of June, sowing 2: 3–4 weeks later) for a total of four environments per year. In each environment, the genotypes were laid out as an augmented block design using the recurrent parents as checks. Because of logistic constraints, all crosses could not be grown in one season. Therefore, some of them were grown in 2012 and others in 2013, each year containing both KK and GR subpopulations (only donor parents were different across years). For each year, only genotypes present in all 4 environments were kept. In 2012, 1821 genotypes from 18 crosses were kept (GR: 13 crosses and 1,403 genotypes, KK: 5 crosses and 418 genotypes), while in 2013, 783 genotypes from 14 crosses were kept (GR: 8 crosses and 528 genotypes, KK: 6 crosses and 255 genotypes). In total, 2498 genotypes from 29 crosses were considered across years (GR: 1848 genotypes, 19 crosses; KK: 650 genotypes, 10 crosses) and only 106 genotypes belonging to three crosses were common to the two years.

Seven traits were measured: flag leaf appearance (FLAG, CO_324:0000631) as the number of days after sowing when half of the plot had their ligulated flag leaves visible, peduncle length (PED, CO_324:0000622) as the distance in cm between the final node and the panicle bottom, panicle length (PAN, CO_324:0000620) as the distance in cm from the end of the peduncle to the panicle top, stem length (STEM, CO_324:0000627) as the distance in cm from the soil to the top of the stem, plant height (PH, CO_324:0000623) as the distance in cm between the soil and the panicle top, number of inter nodes (NODE_N, CO_324:0000605), and yield (YIELD : CO_324:0000403) in kg.ha^-1^ at the plot level. All traits except FLAG were measured at harvest.

### Genotypic data

The 2498 offspring and their parents were genotyped using genotyping by sequencing (GBS, Elshire et al., 2011) with 384-plex libraries on an Illumina HiSeq 2000 sequencer. The offspring were genotyped at generation BC1F3. The sequence data were analysed running the reference genome-based TASSEL GBS pipeline (Glaubitz et al., 2014). Unique tags (3,844,911) were aligned on the sorghum reference genome v2.1 (Paterson et al., 2009). After the filtering of raw genotype data for minor allele frequencies (MAF < 0.05) and single marker missing data (<0.9), 51,545 segregating single nucleotide polymorphisms (SNPs) were identified among the parents with between 11,856 and 26,128 SNPs segregating in the individual crosses. Missing values in the parents were imputed using Beagle (Browning et al., 2018). Missing values in the offspring genotypes were imputed using FSFHap (Bradbury et al., 2007). We determined a unique genetic consensus map by projecting the physical distance of the 51,545 markers on a high-quality genetic consensus map (Guindo et al., 2019) using the R package ziplinR (https://github.com/jframi/ziplinR).

### NIRS data

For each genotype in each year, spectra were acquired in one environment (one site and one sowing date) on sun-dried grain samples after harvest (**Fig. 1**). Spectra were obtained with a MPA Bruker spectrometer (Bruker Optics Inc., Ettlingen, ©) for 1154 wavelengths ranging from 800 to 2800 nm. For each sample, the average of three technical repetitions was used and outlier spectra were removed from the data set afterward.

To assess the effect of spectra pre-processing, smoothing using Savitsky-Golay procedure was applied and four classical chemometric pre-treatments were applied to the smoothed spectra: First derivative (DER1), second derivative (DER2), standard normal variate (SNV), and detrend (DET).

### Estimation of phenotypic breeding values

Due to a lot of missing checks in the design, it was not possible to properly take advantage of the augmented block lay out. Moreover, due to the low number of common genotypes between years, 2012 and 2013 data were analysed separately. For each year, breeding values were estimated with the following model:

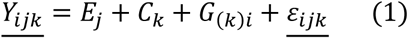

With *Y_ijk_* the phenotypes, *E_j_* the fixed effect of the environment *j* (as a combination of site and sowing date), *C_k_* the fixed effect of the cross 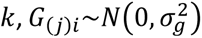 *iid* the random effect of the genotype *i* nested into cross 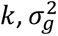 being the global genetic variance across crosses, and *ε_ijk_*~*N*(0, σ^2^) *iid* the residual, σ^2^ being the residual variance. *G*_(_*_k_*_)_*_i_* and *ε_ijk_* were considered independent and the breeding values were estimated as the sum of the cross BLUEs and genotype BLUPs. To minimize the effect of direct relationships between spectra and phenotypes and thus getting closer to an information brought by genotypic data, phenotypic data from the NIRS acquisition environment were not included in breeding values estimations (**Fig. 1**).

### Variance components and heritability

To estimate the heritability of the phenotypic traits, a model similar to model (1) was fitted but considering all terms as random, thus considering 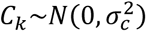 *iid* the random effect of the cross 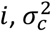 being the variance due to the cross across crosses, 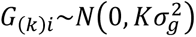 *iid* the random effect of the genotype *i* nested into cross 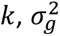 being the genetic variance and *K* being the realized kinship matrix calculated with the Van Raden method (VanRaden, 2008), 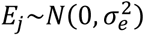 *iid* the random effect of the environment (as a combination of site and sowing date), 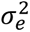 being the environmental variance, and *ε_ijk_*~*N*(0, σ^2^) *iid* the residual, σ^2^ being the residual variance. All effects were considered independent, and heritability was estimated as 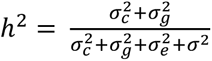. The same model was used to estimate the heritability of each wavelength but without considering an environmental effect and variance as spectra were acquired in only one environment in each year.

### Prediction models

Estimation of prediction accuracies were performed in R (R Core Team, 2023) using 4 different models (**Fig. 1**): GBLUP as a reference with the package sommer (Covarrubias-Pazaran, 2016), partial least square regression (PLSR) with the package rchemo (Brandolini-Bunlon et al., 2023), ridge regression (RIDGE) and least absolute shrinkage and selection operator (LASSO) with the package glmnet (Friedman et al., 2010). GBLUP was used with both genotypic and NIRS data as a reference, while the other models were only used with NIRS data.

### GBLUP model

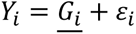

With *Y_i_* the breeding values as calculated in model 1, 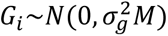 the random effect of genotype 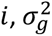 being the genetic variance and *M* being the kinship matrix designated as K when it was based on genotypic information or the pseudo-kinship matrix designated as H when it was based on hyperspectral information, and *ε_i_*~*N*(0, σ^2^) the residual, σ^2^ being the residual variance.

The genotypic kinship matrix *K* was calculated with the Van Raden method (VanRaden, 2008), and the hyperspectral pseudo-kinship matrix *H* was calculated as *H* = *SS^t^*/*n* with *S* being the scaled and centred spectral matrix, *S^t^* being the transpose matrix of *S* and *n* being the number of wavelength (Robert et al., 2022a).

### Ridge and LASSO models

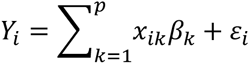

Where the marker effect *β* are estimated by minimizing the following loss function:

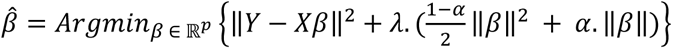

With constraint 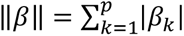 and λ > 0.

With *Y_i_* the breeding values of the genotypes *i* based on model (1), *x_ik_* is the absorbance at wavelength *k* of genotype *i*, and *β_k_* is the effect of the wavelength *k*, and *ε_i_*~*N*(0, σ^2^) *iid* is the residual. LASSO is obtained with *α* = 1 and ridge regression is obtained with *α* = 0. The value of λ was optimized by internal 10-fold cross validation for every model that was computed with built-in functions of the glmnet package.

### PLSR model

PLSR is based on two PCA-like decompositions of the data into latent variables:

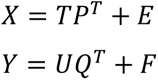

*X* is the *n* × *m* spectral matrix (*n* observations and *m* predictors i.e., wavelengths).

*Y* is the *n* × 1 vector of the response variable (breeding values).

*T* and *U* are *n* × *l* matrices (for *l* latent variables) and are respectively the scores (i.e., projected values of individuals on the latent variables) for *X* and *Y* matrices. They are estimated so that the covariance between column *i* of *U* and column *i* of *T* are maximized while covariance between column *i* of *U* and column *j* of *T* (with *i* ≠ *j*) are zero (*i.e.,* latent variables of T and U are built to be orthogonal).

*P* and *Q* are *m* × *l* matrices and are respectively the loading matrices of *X* and *Y*.

*E* and *F* are respectively the error terms of *X* and *Y* assumed to be independent and identically distributed normal random variables.

Predictions of *Y* are obtained by calculating the scores (*T* matrix) of the validation *X* matrix on the latent variables estimated, then estimating the scores of the unknown *Y* (*U* matrix) matrix through covariances between *U* and *T* matrices and then calculating the estimated values of *Y* through the estimated loadings of *Y* (*Q* matrix).

Predictions were tested with several latent variables ranging from one to 20. Final results were obtained with 10 latent variables as this number maximized PA and minimized Root Mean Square Error of Prediction (RMSEP) overall.

### Cross validation scenarios

To assess the effects of population structure on prediction accuracy, three cross-validation scenarios were used (**Fig. 1**). Those scenarios are comparable to the ones in Lehermeier et al., (2014):

- CV1: intra cross scenario, where a 5-fold cross validation is performed using training and validation set data coming from a single cross
- CV2: intra recurrent parent leave one cross out scenario, where one cross in the validation set is predicted by all the crosses that share the same recurrent parent in the training set
- CV3: inter recurrent parent scenario, where a cross of one population (e.g. GR) is predicted using the crosses of the other population (KK), excluding the crosses with a shared parent

Training set sizes of CV1 were dependant on the specific cross size and ranged from 40 to 108 genotypes. CV2 and CV3 scenarios were also used to assess the effect of training set size on prediction accuracies. In these scenarios, different training set sizes were set by increasing the number of genotypes from 50 to 1250 with a step of 50. For each training set size, sampling was made so that the structure of the population was respected (i.e., the proportion of genotypes belonging to each cross was the same as in the total population in all training sets). To study the other impacting factors, CV2 scenario was used with training sets sizes of 1000 genotypes in 2012 to have a large training set and keep a reasonable computing time, and 450 in 2013 to have the largest possible training set given the lower number of genotypes in 2013. For each scenario and different population size within scenario (CV2 and CV3), 10 replications were carried out.

An additional scenario (CV4) was developed to study the impact of NIRS acquisition environment (**Fig. 1**): inter-year scenario with crosses from one year predicted by a model trained on crosses from the other year so that spectra from the training and validation set were acquired in different environments and different genotypes. CV4 was trained with data collected in 2012 and validated with data from 2013 (CV4_2013) and vice versa (CV4_2012). Even though training and validation sets share no genotypes, crosses from both recurrent parents are present in both sets. For this scenarios, 30 repetitions were made, and training set size was set to 600, respecting the structure of the population, to have the biggest training set size possible given the number of genotypes in 2013. **Table 1** is a summary of scenarios and training set sizes used to study each factor.

### CDmean and Fst calculation

To quantify relatedness between training and validation sets in CV1, CV2, and CV3, CDmean and Fst indicators were calculated. Fst were calculated with the R package BEDASSLE (Bradburd, 2024) which estimates Fst according to Weir and Hill, (2002). CDmeans were calculated according to Rincent et al., (2012) with R codes given in the TrainSel package (Akdemir et al., 2021) documentation. Then, linear regressions between PA and Fst or CDmean were carried out, and the effects of these indicators on PA were tested with a Student test on the slopes of the regressions.

### Wavelength selection

To study the effect of wavelength selection, three steps were implemented. First, LASSO and PLSR models were used to estimate the importance of each wavelength in prediction with the loadings of the PLSR and the weights of the LASSO. Then, wavelengths were selected based on these loadings and weights. Finally, the selected wavelengths were used in GBLUP models to compare a GBLUP with all wavelengths and a GBLUP with only selected wavelengths. Training and validation sets were not the same in the LASSO/PLSR and in the GBLUP model to avoid overfitting. PLSR and LASSO models were trained with CV2 training sets with 2012 and 2013 data with 450 genotypes, which correspond to the largest training set that could be composed using 2013 data. According to these analyses, the loadings of the first latent variable of PLSR and the weights attributed to each wavelength in LASSO were retrieved. The mean loadings and weights were calculated for each year. The selection of wavelength was based on the local maxima and minima of the mean loadings and maxima of the weights. The selected wavelengths were then used in a GBLUP model in CV2 scenario with training set sizes of 1000 for 2012 data and 450 for 2013 data. See Fig. S1 for a schematic representation.

Another method of wavelength selection was random selection. In this method, 10 samples from 1 to 10, twenty, thirty, forty, fifty and sixty wavelengths were randomly chosen to calculate the kinship matrix. Each sample was then used in predictions using the CV2 scenario with training set sizes of 1000 for 2012 data and 450 with 2013 data.

All analysis related to wavelength selection were performed on raw spectra to mimic at best what could be achieved with cheaper sensors measuring few wavelengths as pre-processing the same wavelength with 1154 or less than 60 wavelengths would not yield the same spectral values.

### Models’ accuracies metrics

The PA of all models were calculated with the Pearson’s correlation coefficient between observed and predicted breeding values. As CV scenarios were based on a leave one cross out (or alike) methods, PAs were always calculated within cross for each scenario. Thus, the results correspond to the within cross PA metrics average over the different crosses. Significant differences between PA were tested using pairwise Wilcoxon tests with a Bonferroni correction if not mentioned otherwise.

### Graphics

All graphics were produced with the ggplot2 package (Wickham, 2016) of R (R Core Team, 2023), or with PowerPoint.

## Results

### Description of phenotypic data

Coefficients of variation for each trait in each environment are available in Table S1. The trait with the highest variability was YIELD, followed by PED, STEM and PH, followed by NIN and PAN, followed by FLAG which was the least variable trait. For PAN, PED, PH, FLAG and NIN, coefficients of variation were stable across environments and years. STEM was more variable for 2012 than 2013 overall. YIELD was more variable in 2013 than 2012 overall except for SB1 in 2012 where the highest coefficient of variation (0.7) occurred.

Correlation between environments for each trait are presented in Fig. S2. PED was the trait with the highest correlation between environments ranging from 0.62 to 0.83. YIELD was the trait with the lowest correlations between environments, ranging from 0.06 to 0.39. FLAG, PH and STEM had medium to high correlations between environments, ranging from 0.52 to 0.83. NIN had low to medium correlation between environment, ranging from 0.2 to 0.47. Finally, PAN had medium-high correlation between environment in 2013 ranging from 0.65 to 0.67, and medium-low correlations in 2012, ranging from 0.39 to 0.52.

### Variance partitions of phenotypic traits and spectra

Partition of variance for each trait are presented in **Fig. 2**. Heritability estimates were generally lower for 2013 data. Traits related to plant height (PED, PH, and STEM) were the most heritable in both years with heritability ranging between 0.46 and 0.48 in 2012 and between 0.28 and 0.33 in 2013. YIELD and NIN were the least heritable traits in both years with heritability ranging between 0.11 and 0.18. FLAG had intermediate heritability with values of 0.29 and 0.26 in 2012 and 2013, respectively. PAN had among the lowest heritability in 2012 with a value of 0.22 while it was close to the highest in 2013 with a value of 0.28. Heritability of the wavelengths appeared to be stable along the spectra with a severe drop around 2700 nm. Mean heritability in 2012 was 0.51, and mean heritability in 2013 was 0.44. Overall, the variance due to the cross was very low, which may be due to the use of a marker derived kinship that probably capture both the population structure represented by the cross and within cross individual differences.

**Fig. 2.**
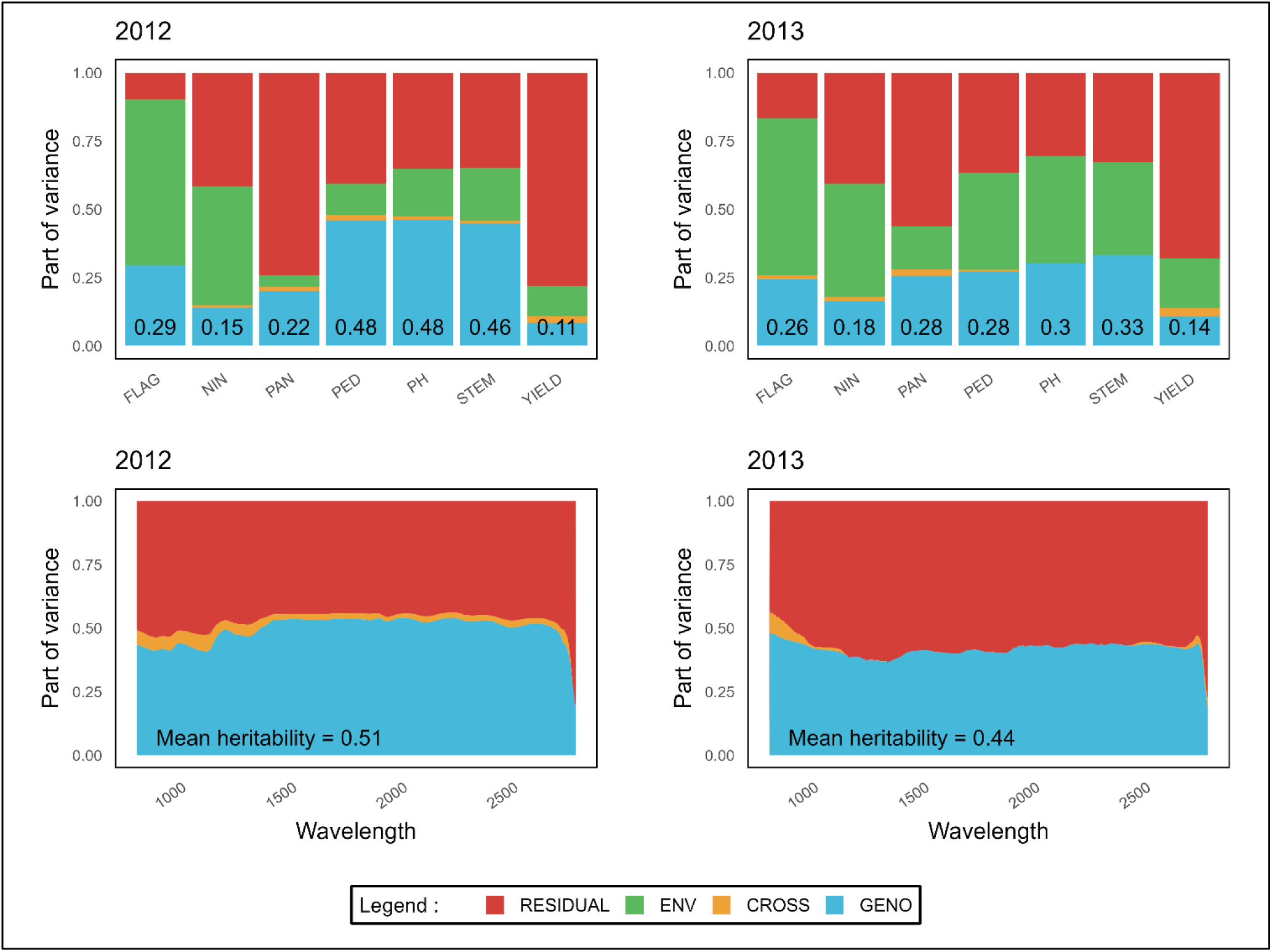
Partition of variance for the studied traits and wavelengths along spectra in 2012 and 2013. RESIDUAL, ENV, CROSS and GENO stand for the factors contributing to the phenotypic variance. Numbers in black are heritability of traits.

### Overall comparison of GP and PP

**Fig. 3** presents PAs of genomic prediction and phenomic prediction for raw and pre-processed spectra in the CV2 scenario with 1000 genotypes in the training set in 2012 and 450 in 2013. In 2012, FLAG and NIN, PP and GP performed similarly. Only PP with spectra pre-processed with DER2 and SNV had significantly lower PAs than GP for NIN. For plant height related traits (PED, PH, STEM), and PAN, PP systematically had lower accuracy than GP. For YIELD, PAs of PP were significantly lower than PAs of GP but differences in PAs median were not as large as for plant height related traits.

**Fig. 3.**
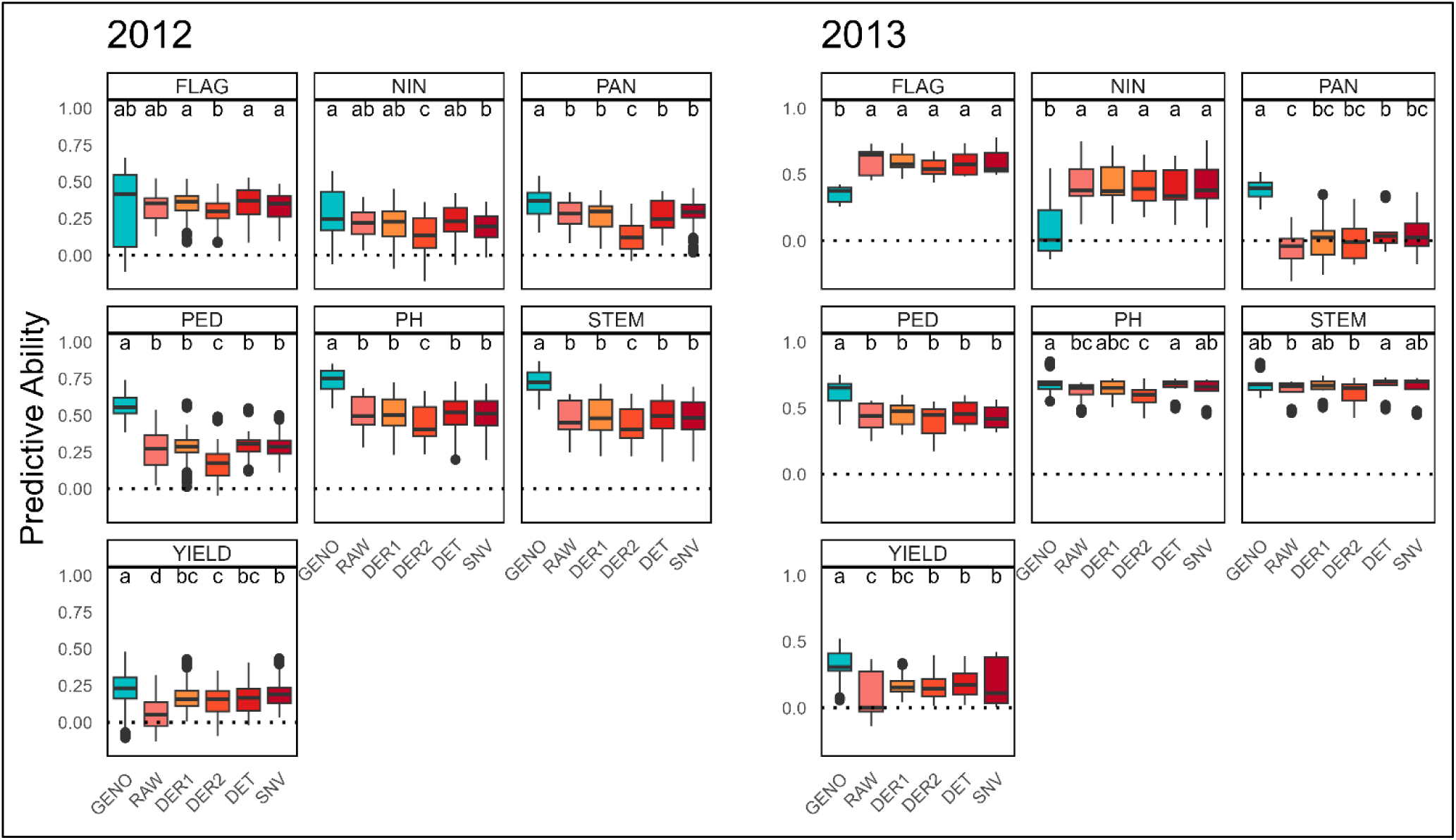
Effects of spectra pre-processing on phenomic prediction predictive abilities. Predictive abilities were calculated using GBLUP in the CV2 scenario with a training set size of 1000 in 2012 and 450 in 2013. Letters represent the results of pairwise Wilcoxon tests corrected by Bonferroni method for a threshold of 5%. GENO = genomic prediction, RAW = phenomic prediction with raw spectra, DER1 and DER2 = phenomic prediction with first and second derivative of the spectra, DET = phenomic prediction with spectra processed with detrend, SNV = phenomic prediction with spectra processed with the standard normal variate method. Horizontal dotted line is at the 0 value. FLAG = flag leaf appearance, NIN = number of inter-nodes, PAN = panicle length, PED = peduncle length, PH = plant height, STEM = stem length, YIELD = yield.

Slightly contrasting results were observed in 2013. In this year, PP and GP performed similarly for PH and STEM. PP consistently outperformed GP for FLAG and NIN, but GP consistently outperformed PP for YIELD. In addition, PP was not able to predict PAN for this year.

### Effect of spectra pre-processing

In 2012 (**Fig. 3**), for all trait except YIELD, only DER2 lead to significantly lower PAs while all other pre-processing did not lead to significant differences in PAs. In 2013, no significant differences between pre-processing were found for FLAG, NIN, and PED. Some significant differences were found between DER2 and DET for STEM. For PH, DER2 gave significantly lower PAs than DET and SNV, and RAW spectra gave significantly lower PAs than DET. Interestingly, all pre-processing increased the PAs for YIELD in 2012 and 2013 with no significant differences between them except for DER2 which had significantly lower results than SNV in 2012. As DET, DER1 and SNV gave the best performances overall with no significant differences, and SNV had the highest PAs for YIELD in 2012, following results were computed using SNV pre-processed spectra.

### Effect of statistical method

**Fig. 4** presents PAs for PP obtained from 4 statistical models: GBLUP, ridge, LASSO and PLSR. PAs were obtained in the CV2 scenario with a training set size of 1000 genotypes in 2012 and 450 in 2013. In 2012, no effect of statistical method was observed for NIN, PAN, PH, and STEM. Significant differences were found between methods for FLAG, PED and YIELD but they were inconsistent across traits. The highest difference was found between PLSR and LASSO for FLAG, being 0.09, but all other significant differences that were found represent a gap smaller than 0.05. With 2013 data, significant differences were only found for PED, where Ridge and LASSO had PAs of 0.54 and 0.52 against of 0.42 for GBLUP. Overall, no statistical model had a clear advantage over the others. Next analyses were carried out using the GBLUP reference model.

**Fig. 4.**
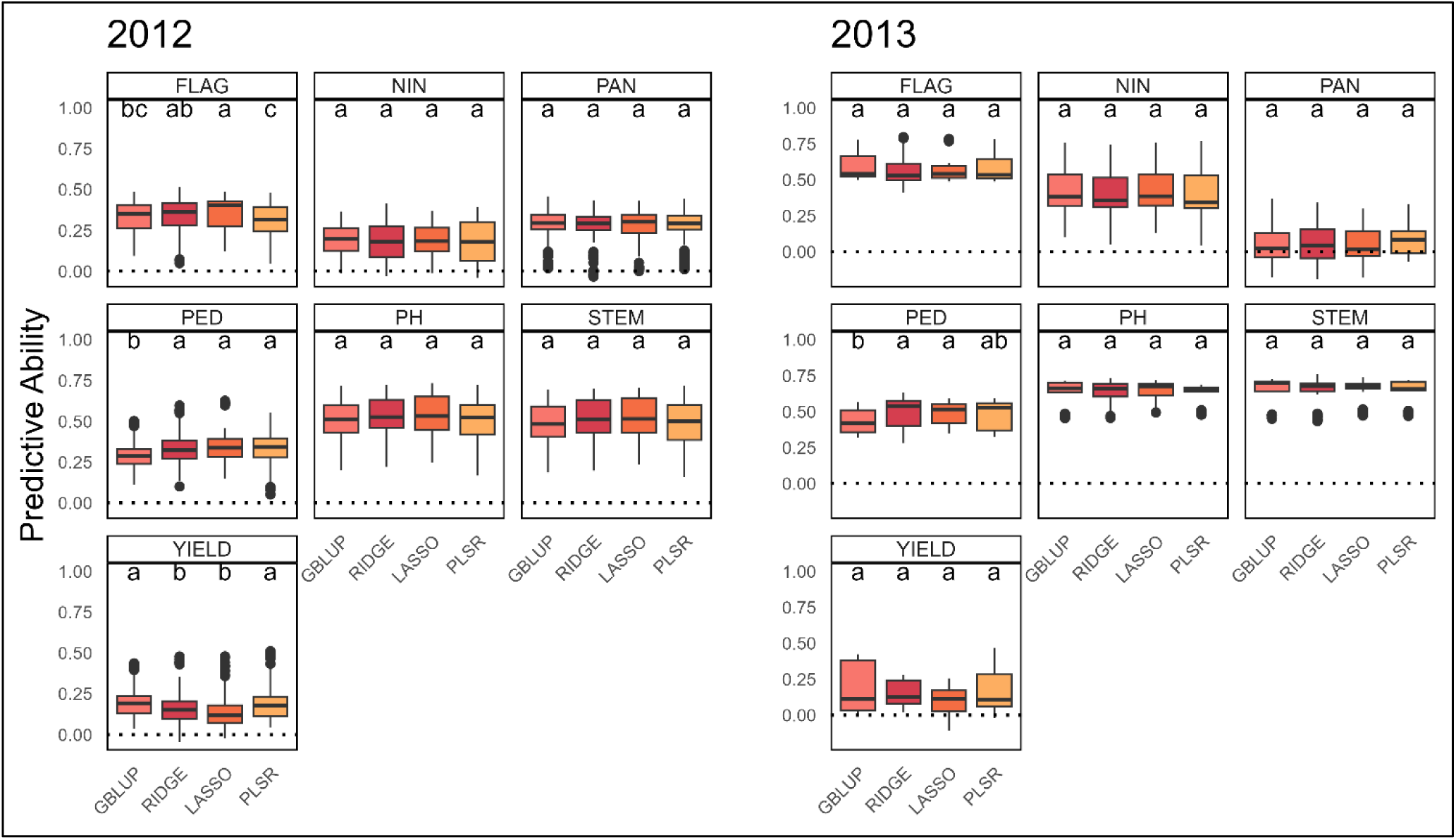
Effects of statistical method on predictive abilities. Predictive abilities were calculated in the CV2 scenario with a training set size of 1000 for 2012 and 450 for 2013, and spectra pre-processed with SNV. Letters represent the results of pairwise Wilcoxon tests corrected by Bonferroni method for a threshold of 5%. Horizontal dotted line is at the 0 value. FLAG = flag leaf appearance, NIN = number of inter-nodes, PAN = panicle length, PED = peduncle length, PH = plant height, STEM = stem length, YIELD = yield. GBLUP = reference gblup model, RIDGE = ridge regression, LASSO = least absolute selection and shrinkage operator, PLSR = partial least square regression.

### Effect of training set size

The evolution of PAs according to the training set size for the CV2 scenario in 2012 using a GBLUP model is presented in **Fig. 5**. For FLAG, NIN and PAN, PP reached a plateau of PAs around 500 genotypes in the training set while PAs of GP continued increasing. For NIN and PAN, PAs of GP ended up higher than PAs of PP but for FLAG, PAs of GP ended up at the same level as PAs of PP. For plant height related traits, the evolution of PP and GP had a similar shape but with an offset, PAs of PP being lower than PAs of GP. For YIELD, GP PAs increased faster than PP PAs, but both prediction methods reached the same levels with large training set sizes. The effect of training set size was studied with 2013 data (not shown) but to the lower number of genotypes available (maximum training set size of 450) was not enough to draw conclusions. The effect of training set size was also studied with CV3 scenario (Fig. S3). PAs had a much greater variability for each training set size making results hardly interpretable, and no pattern emerges from these analyses.

**Fig. 5.**
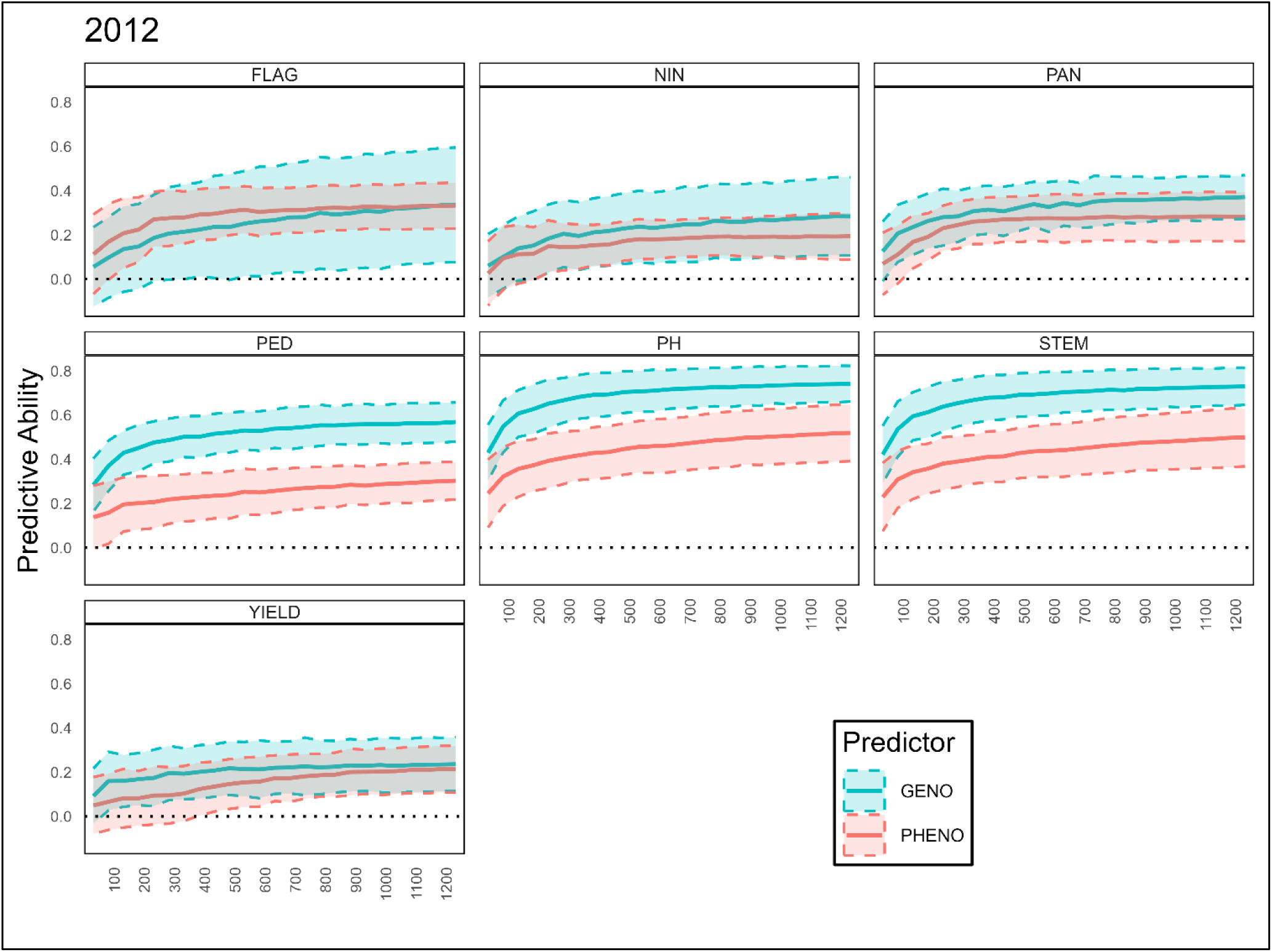
Effect of training set size on predictive abilities. Predictive abilities were calculated with 2012 data, using GBLUP in the CV2 scenario and spectra pre-processed with SNV. Horizontal dotted line is at the 0 value. FLAG = flag leaf appearance, NIN = number of inter-nodes, PAN = panicle length, PED = peduncle length, PH = plant height, STEM = stem length, YIELD = yield. Full lines represent the mean PAs and dotted lines represent the standard error around the mean PA.

### Effects of population structure

**Fig. 6** presents the PAs for CV1, CV2 and CV3 scenarios for a training set size of 100 (which is the maximum training set size available in CV1) using a GBLUP model on 2012 data. The BCNAM population is strongly structured as can be seen by a PCA on SNP or spectral data (Fig. S4). Thus, it is relevant to build prediction scenarios with varying relatedness between training and validation sets. This relatedness decreased from CV1 to CV3 (Fig. S5) which caused PAs to decrease both in PP and GP (**Fig. 6**). GP had more significant differences in PA between the CV scenarios than PP. It was especially the case for plant height related traits. For instance, the median PAs of PP for PH dropped from 0.44 to 0.21 between CV1 and CV3, while for GP it dropped from 0.72 to 0.08. For all traits except NIN and PAN, PP had significantly higher PAs than GP in CV3 scenario while GP had significantly superior PAs in CV1 and CV2. For NIN and PAN, PAs of PP were not significantly different from PAs of GP in CV3, but were already close to 0 in CV2, hindering a relevant comparison concerning the stability of PAs across scenarios.

**Fig. 6.**
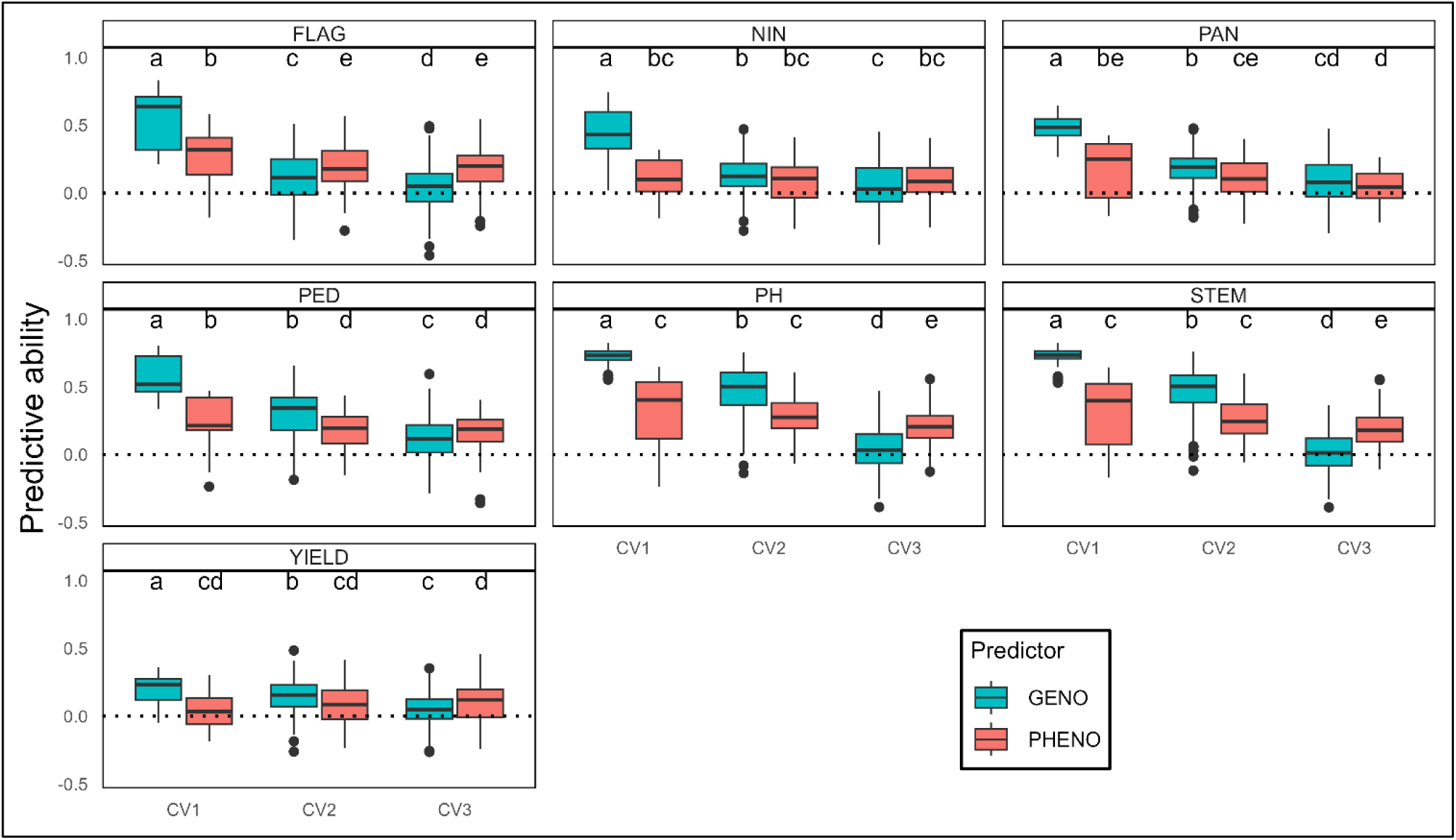
Effects of population structure on predictive abilities. Predictive abilities were calculated with 2012 data, using GBLUP scenario with a training set size of 100 and spectra pre-processed with SNV. Horizontal dotted line is at the 0 value. Letters represent the results of pairwise Wilcoxon tests corrected by Bonferroni method for a threshold of 5%. FLAG = flag leaf appearance, NIN = number of inter-nodes, PAN = panicle length, PED = peduncle length, PH = plant height, STEM = stem length, YIELD = yield. CV1, CV2, CV3 refer to cross-validation scenarios with decreasing genetic relatedness between training and validation sets from CV1 to CV3. CV1 = intra-cross 5-fold cross validation, CV2 = intra recurrent parent leave one cross out cross-validation, CV3 = inter recurrent parent cross-validation.

CDmean and Fst indicators were calculated for every cross-validation partition used to study population structure and linear regressions were carried out to test the effect of CDmean and Fst between training and validation sets on PA (Fig. S6). Slopes for PP were systematically closer to 0 than slopes for GP. Some slopes for PP were not even significatively different from 0: NIN, PH, STEM and YIELD for CDmean, and NIN for Fst. There are no CDmean values between 0.32 and 0.56, and no Fst values between 0.11 and 0.51 in our partitions. Optimizing training and validation set composition to target specific CDmean or Fst values is extremely time consuming. Given the variability of PA for each CDmean or Fst value, we did not seek to optimize partitions to access intermediate values because it would have taken too much time to have a representative number of partitions.

The effect of population structure was not studied in 2013 because the maximum training set size for CV1 scenario was too low to achieve relevant PA.

### Effect of NIRS acquisition environment

Results for inter-year prediction are presented in **Fig. 7**. CV2_2012 is the reference scenario with data of the training and validation set coming from 2012, and predicted with the CV2 scenario. In CV4_2012, spectra and breeding values used to train the model were acquired in 2013 while spectra and breeding values used to validate the model were acquired in 2012. In CV4_2013, training data were acquired in 2012 and validation data were acquired in 2013. Training and validation sets of CV4 scenarios share no common environment nor common genotypes. For plant height related traits, results of GP were more stable across scenarios than results of PP. For FLAG and YIELD, PP and GP performed similarly across scenarios. For NIN, GP and PP were both stable across scenarios but PP had lower PAs. For PAN, PP had no predictive power in the inter-year scenario which is not surprising given the results of PA for this trait with 2013 data only (**Fig. 3**). For FLAG, PAN, PED, PH and STEM, training a model on 2012 data gave significantly higher PAs than training models on 2013 data, and the opposite occurred for YIELD.

**Fig. 7.**
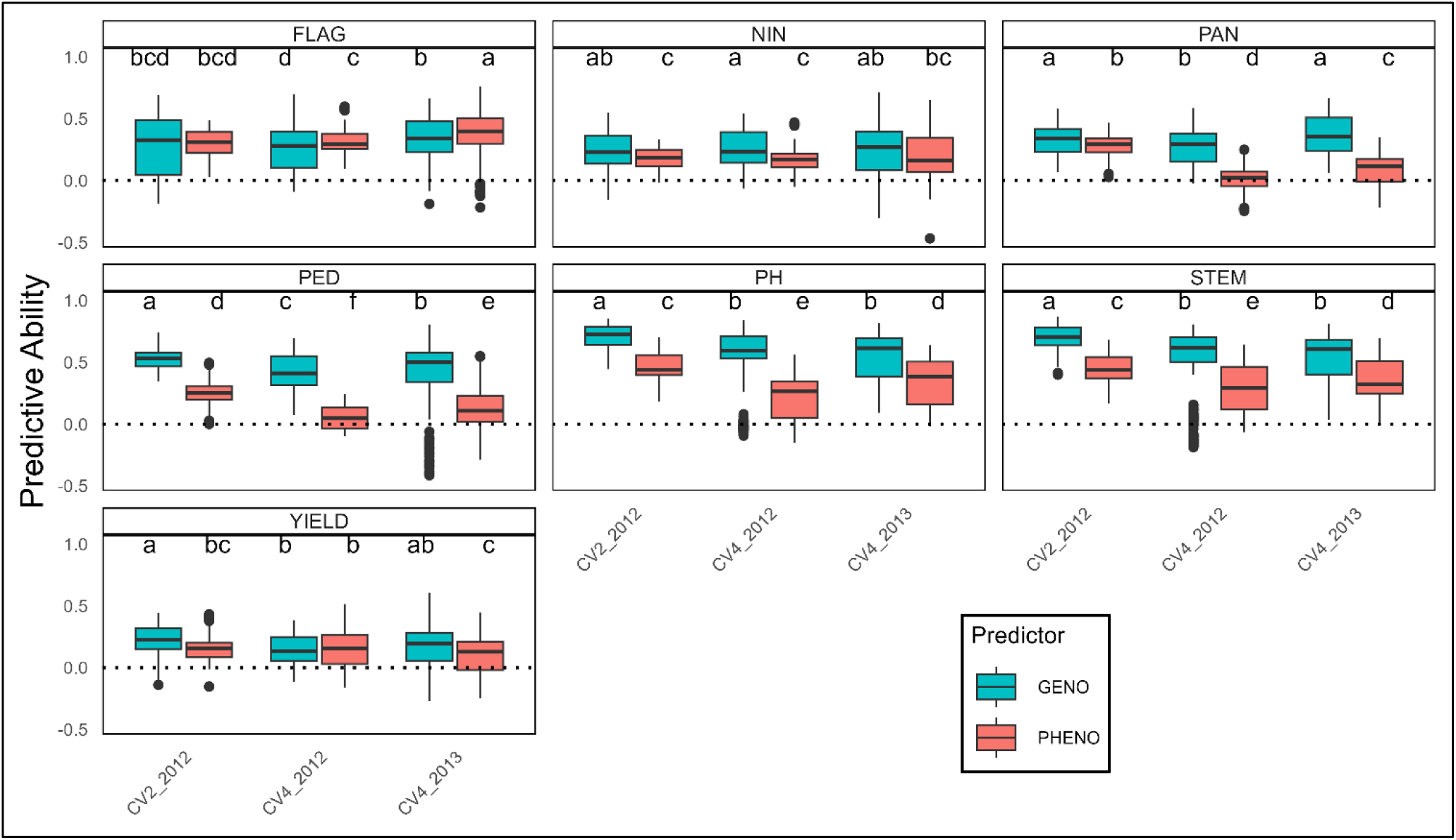
Effects of NIRS acquisition environment on predictive abilities. Predictive abilities were calculated using GBLUP with training set size of 600 and spectra pre-processed with SNV. CV2_2012 is the reference model, in CV2 scenario for 2012 data with 600 individuals in the training set and spectra pre-processed with SNV. CV4_2012 = model trained on 2013 data and validated on 2012 data, CV4_2013 = model trained on 2012 data and validated on 2013 data. Horizontal dotted line is at the 0 value. Letters represent the results of pairwise Wilcoxon tests corrected by Bonferroni method for a threshold of 5%. FLAG = flag leaf appearance, NIN = number of inter-nodes, PAN = panicle length, PED = peduncle length, PH = plant height, STEM = stem length, YIELD = yield.

### Effect of wavelength selection

The results of PP using wavelengths selected by PLS, LASSO, or randomly are presented in **Fig. 8**. Models with wavelength selection were compared to PP models with all wavelengths and GP. The wavelengths that were selected by LASSO or PLSR are presented in Fig. S7, S8, S9 and S10. The sets of wavelengths selected by LASSO gave the same PAs as using all wavelengths for all trait in 2012 and 2013. Except for FLAG, PED, and NIN, wavelengths selected by PLS gave lower PAs than PP based on all wavelengths in 2012. In 2013, PLS selected wavelengths gave similar PAs as PP based on all wavelength except for plant height related traits.

**Fig. 8.**
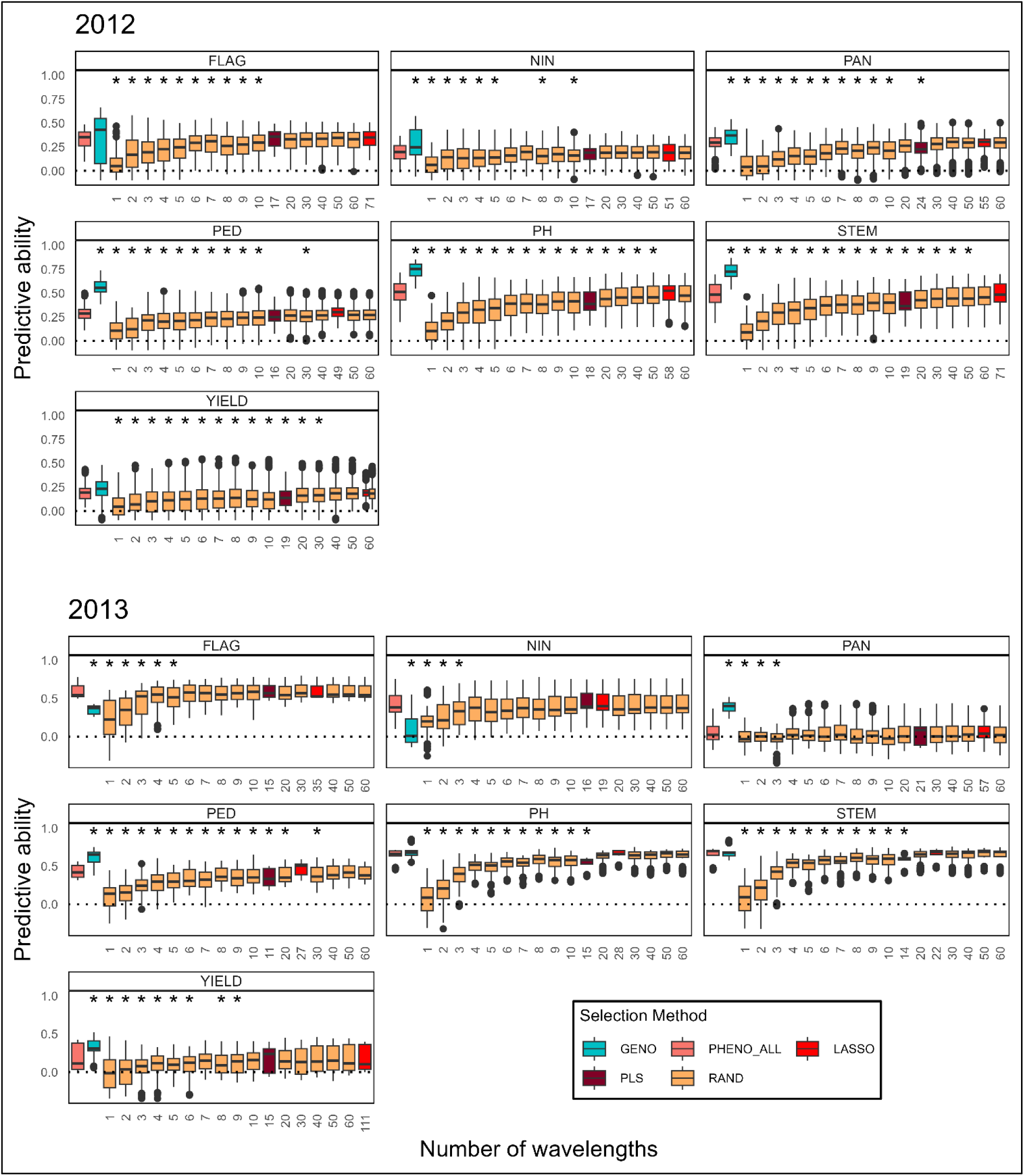
Effects of wavelength selection on predictive abilities. Predictive abilities were using GBLUP in the CV2 scenario with a training set size of 1000 in 2012 and 450 in 2013, and spectra pre-processed with SNV. Wavelengths were randomly selected (RAND), or selected using LASSO and PLS models. PHENO_ALL is the phenomic prediction with all wavelengths, and GENO is the genomic prediction. Horizontal dotted line is at the 0 value. Asterisks indicate predictive abilities that are significantly different from the PHENO_ALL modality, tested with a Dunnett test with a threshold of 5%. FLAG = flag leaf appearance, NIN = number of inter-nodes, PAN = panicle length, PED = peduncle length, PH = plant height, STEM = stem length, YIELD = yield. The x-axes are not linear and differ across traits and years as different number of wavelengths were selected by PLS and LASSO in these different contexts.

Considering randomly selected wavelengths, the number of wavelengths required to reach the same PAs as using all wavelengths differed across traits. In 2012, slightly more than 10 wavelengths were required to predict FLAG NIN, PAN and PED with the same PA as with all wavelengths. However, more than 30 were needed for YIELD, while more than 50 wavelengths were needed for PH and STEM. In 2013, only 4 wavelengths were sufficient to predict NIN, while 6 were needed for FLAG, more than 10 for YIELD, PH, and STEM, and more than 30 for PED. For equivalent number of wavelengths selected, LASSO and PLS methods sometime have a slight advantage in median PA, and less variability for plant height related traits. However, LASSO and PLS methods did not generally outperform random selection. Lastly, randomly selecting a very low number of wavelengths gave significantly lower PAs than using all the wavelengths, yet, the predictive power reached at only 3 randomly selected wavelength was surprisingly high.

## Discussion

In this study, we implemented PP on sorghum for the first time using a large BCNAM population. Our design comprising 2 years with 4 environments each, and a strongly structured population allowed us to study factors impacting PP, namely, spectra pre-processing, statistical method, training set size, population structure, NIRS acquisition environment, and wavelength selection.

### Phenomic selection for sorghum breeding

Despite being the fifth most produced cereal worldwide, a major crop for food security in arid and semi-arid areas, and a promising crop to face climate change issues, sorghum genetics and breeding programs have not received much attention and means for development and optimization (Khoury et al., 2014; Pingali and Traxler, 2002). GP has already been tested on this species with promising results in temperate programs (Hunt et al., 2018; Maulana et al., 2023; Yu et al., 2016) which already use spectral data to evaluated forage and grain quality. Moreover, more and more sorghum breeding programs from temperate and semi-arid regions are mobilizing NIRS technologies on a routine basis to assess grain and forage quality. Thus, the additional costs to implement PP in these programs is low, and PP fits well in the current context of sorghum breeding.

More generally, PP as a low-cost, or even low-tech method is especially relevant for breeding programs with low financial support and access to molecular technologies. Our study demonstrated the possibility of using PP even in a rather unfavourable very low cost set up *i.e.*, with only one reference NIRS acquisition environment and one spectrum per genotype. No repetition of spectral measurement makes it likely that genetic information is hard to retrieve, but still, models were able to grasp information out of them. Moreover, we showed that using a single reference NIRS acquisition environment is possible as shown by previous studies (Brault et al., 2022; Rincent et al., 2018; Zhu et al., 2022, 2021). We also showed that predicting untested genotypes in untested environment is possible even though PP often yields lower PAs than GP in this case. One interesting approach to further evaluate the possibilities of using PP in such unfavourable design would be to better characterize environments or quantify environmental relatedness between training and validation sets to see how it affects PA. Such approach currently applied for GP yielded significant improvements (Montesinos-López et al., 2024).

### Factors impacting PP

Our results show that PP can impact the choice of the training set. The results on FLAG, NIN and PAN are in accordance with previous studies showing that PP needs smaller training sets than GP to reach a plateau of PAs (Dallinger et al., 2023; Zhu et al., 2022, 2021). Contrarily to our results, Zhu et al., (2021) showed that this feature can also be true for plant height and yield. On the other hand, (Zhu et al., 2022), showed that PP sensibility to training set size depends on the breeding population, which may explain this difference.

Results on plant height related traits and FLAG are in accordance with previous studies showing that PP is less affected by population structure (Laurençon et al., 2024; Roscher-Ehrig et al., 2024; Weiß et al., 2022; Zhu et al., 2022, 2021). Most importantly, PP had higher PA than GP for almost all traits in the scenario with the least relatedness between training and validation sets, highlighting a real advantage of PP over GP in less favourable prediction scenarios. Nevertheless, as training set sizes available for comparison between all CV scenarios were small, it would be interesting to test similar scenarios with larger training set sizes to generalize results. This study also highlighted the trait dependency of these features of PP mitigating the advantages PP can have over GP. This trait dependency could not be explained by the genetic architecture. Indeed, no pattern emerged when trying to establish a link between our prediction results and the genetic architecture of traits previously studied in the same population (Garin et al., 2024). Few authors studied the impact of genetic architecture on PP (Roscher-Ehrig et al., 2024; Zhu et al., 2022). They found that PP performs better than GP for complex traits determined by many QTL. The absence of such pattern in our results may arise from the fact that the number of QTL detected for each trait in our population (Garin et al., 2024) is always inferior to 6 and that the explained variance varies a lot for each trait across environments and sub-populations.

Considering factors related to data processing, we found small or no effects of spectra pre-processing and poor performances when spectra were pre-processed with second derivative. This result is hardly generalisable as no consensus exists in the literature regarding the best pre-processing method. Even though some studies found no effect of spectra pre-processing with classical chemometric methods (Brault et al., 2022; Dallinger et al., 2023), it was proposed by Meyenberg et al., (2024) that fine tuning parameters of pre-processing methods could have an impact on prediction. Some improvement of PP performance was found by using genetic values of reflectance or absorbances at each wavelength (Brault et al., 2022). Unfortunately, spectra of our data set were only acquired in one environment per genotype hindering the calculation of genetic values at each wavelength without using a marker-based kinship matrix. While this approach could have been interesting to test, it would have been more a particular case of integrating genomic and phenomic data than a kind of pre-processing that would involve only spectral information.

Finally, considering factors related to data analysis, we did not find any strong effect of the statistical method on the performance of PP. One limit of our study is that we have only tested frequentist linear methods. Regarding this factor, it is also hard to generalise results as some studies including Bayesian and non-linear frameworks did not find major differences between statistical methods (Roscher-Ehrig et al., 2024; Zhu et al., 2021) while others found that using non-linear methods (Cuevas et al., 2019) or deep learning (Mora-Poblete et al., 2024) could enhance PA of PP. Nonetheless, using Bayesian or deep learning methods requires more computational resources than classic frequentist methods and in addition implementing deep learning strategies requires expertise and energy demanding strategies to fine tune model parameters (Lee et al., 2022; Zhang et al., 2019). In this context, applications of such methodologies will be dependent on the effective gains they will provide to compensate their higher investments requirements.

### Taking advantage of the integrative properties of PP to further reduce its implementation costs

Reducing the number of wavelengths used to estimate the kinship matrix could be a way to further reduce the costs of PP as it would allow to use cheaper material acquiring few wavelengths. Selecting wavelengths with PLSR or LASSO models worked well to reduce the number of wavelengths without substantial loss of PA. These approaches could be limited as the wavelengths selected are trait and environment specific, meaning that we could not use a set of selected wavelengths in all cases which is the way to reduce the costs. However, our results with random selection of wavelengths show that this limit can be overcome and that a wavelength selection method is not really required. Indeed, randomly reducing the number of wavelengths to only 10 (with only one acquisition time and no repetitions of genotypes) did not decrease much PA in most cases, meaning that low-cost camera using only a few wavelengths could be used. This result is in line with the “minimal setup” proposed by Mróz et al., (2024) where only three bands in the visible range acquired on a single date for canopy measurement could predict wheat grain yield with an accuracy ranging from 0.51 to 0.55. However, randomly selecting wavelengths could make predictions less stable than selecting wavelengths with non-random methods, which may make the use of regular infrared spectroscopy preferable when affordable.

In this study, PP was used as defined by Rincent et al., (2018), *i.e.*, using near infra-red spectroscopy on plant tissue to estimate a kinship matrix and perform predictions. Since 2018, many studies have widened the concept and have also used canopy measurement to capture fewer visible and NIRS bands with multi-spectral cameras (Fumia et al., 2023b; Galán et al., 2020; Krause et al., 2019), sometimes capturing less than 10 bands (Li et al., 2023; Maggiorelli et al., 2024; Mróz et al., 2024). Other studies used vegetation indexes also calculated with very few bands but often densely measured throughout the growing cycle and coupled with plant height measurements (Adak et al., 2024, 2023b, 2023a, 2022; Biswas et al., 2021; Kaushal et al., 2024; Sandhu et al., 2021; Shafiee et al., 2023; Togninalli et al., 2023; Washburn et al., 2024; Winn et al., 2023), or even temperature and canopy area (Parmley et al., 2019). It would be interesting to study the costs and benefits of using only a few bands in a minimal setup or with dense acquisition along the plant cycle, compared to PP as defined in Rincent et al., (2018).

We hypothesised that using randomly selected wavelengths is possible either because of a high redundancy in spectral information or because of poor use of complete spectral information. The high redundancy hypothesis is based on a linkage-disequilibrium-like relation between uninformative and informative wavelengths due to high collinearity between adjacent wavelengths. Considering this feature of spectra, it is not surprising that a random selection of wavelengths can maintain PA especially when randomly selecting in such a wide range of wavelengths which gives a high chance of selecting different uncorrelated wavelengths each of which are correlated to a different informative one. However, it is interesting that selecting only a few bands (*i.e.*, less than 10) can maintain PA. The poor use of complete spectral information hypothesis is that models do not really extract all the information possible when all wavelengths are used and in fact extract a quantity of information equivalent to what is contained in only a few wavelengths. This could be because wavelengths are considered individually and independently in models (*e.g.*, when calculating a spectral kinship matrix) while they actually are very related to each other. More classical uses of NIRS made in the field of chemometrics rather calculate “features” of spectra (*e.g.*, principal components) to further analyse them (Jean-Michel Roger, personal communication, January 7, 2025).

The possibility to use few wavelengths questions the need for “omic” information in prediction, which are typically quite costly. As stated in Zhu et al., (2022, 2021), each wavelength, behaving like endophenotypes may contain a lot of information including additive, non-additive, environmental and genotype-environment-interaction effects. As such, using a reduced number of bands could in fact contain an omic type of information despite not being as dense and heavy as classic omics.

That being said, all the non-additive information supposedly contained in the spectra were not explicitly taken into account in our models, and it is impossible at this stage to confirm that this information is responsible for PP performances in this case. The growing body of results concerning PP that could be explained by the endophenotype behaviour hypothesis calls for a better understanding of the information contained in the spectra and how it can be linked to genetic relationship matrices. It would be interesting to decompose the phenomic signal into its additive and non-additive parts to compare models that contain both or not. Further analysis could also focus on an efficient way to make this decomposition, which may borrow from the chemometrics framework.

## Conclusion and perspectives

The interest of PP for sorghum breeding has been demonstrated. We studied factors impacting PP, confirmed some of its advantages over GP concerning training population definition and size, but we also highlighted a strong trait dependency of these advantages. We also stressed the robustness of PP in unfavourable application conditions, and when using only a few wavelengths, opening prospects for the adoption of PP in breeding programs from the global south. Further studies are still needed to better understand and decompose phenomic information to use each component at best in predictive models.

## Statement and declarations

### Funding

This work was supported by a grant from the Generation Challenge Programme (Project Numbers G7010.05.01 and G7010.05.02). The work of Clément Bienvenu was supported by a doctoral allowance from the French ministry of higher education and research.

### Competing interest

The authors have no relevant financial or non-financial interests to disclose.

### Authors contribution

KT, MLT, CD, JFR and MV conducted population development, field experiments, field phenotypic data collection and sample preparation. MCS performed the acquisition of NIRS spectra. AB, CC, MV, FdB genotyped the population. CB led the data analysis and prepared the first draft of the manuscript. VG, NS, VS, DP and HdV contributed to the data analysis. HdV, DP and VS supervised the overall research and edited the manuscript. HdV, JFR, DP and VS conceptualized, supervised, financially supported the overall research, and edited the manuscript. All authors revised and approved the manuscript for submission.

### Data availability

All data used in this study is available at https://doi.org/10.18167/DVN1/R70XDD

## Acknowledgement

We acknowledge the MGX-Genotyping platform (UMR AGAP - CIRAD, Montpellier, France) for its support regarding the genotyping of the BCNAM population

## Notes

https://doi.org/10.18167/DVN1/R70XDD

### Competing Interest Statement

The authors have declared no competing interest.

