## Supplementary Materials for "Factors Influencing Phenomic Prediction: A Case Study on a Large Sorghum BCNAM Population"

Journal : Theoretical and Applied Genetics

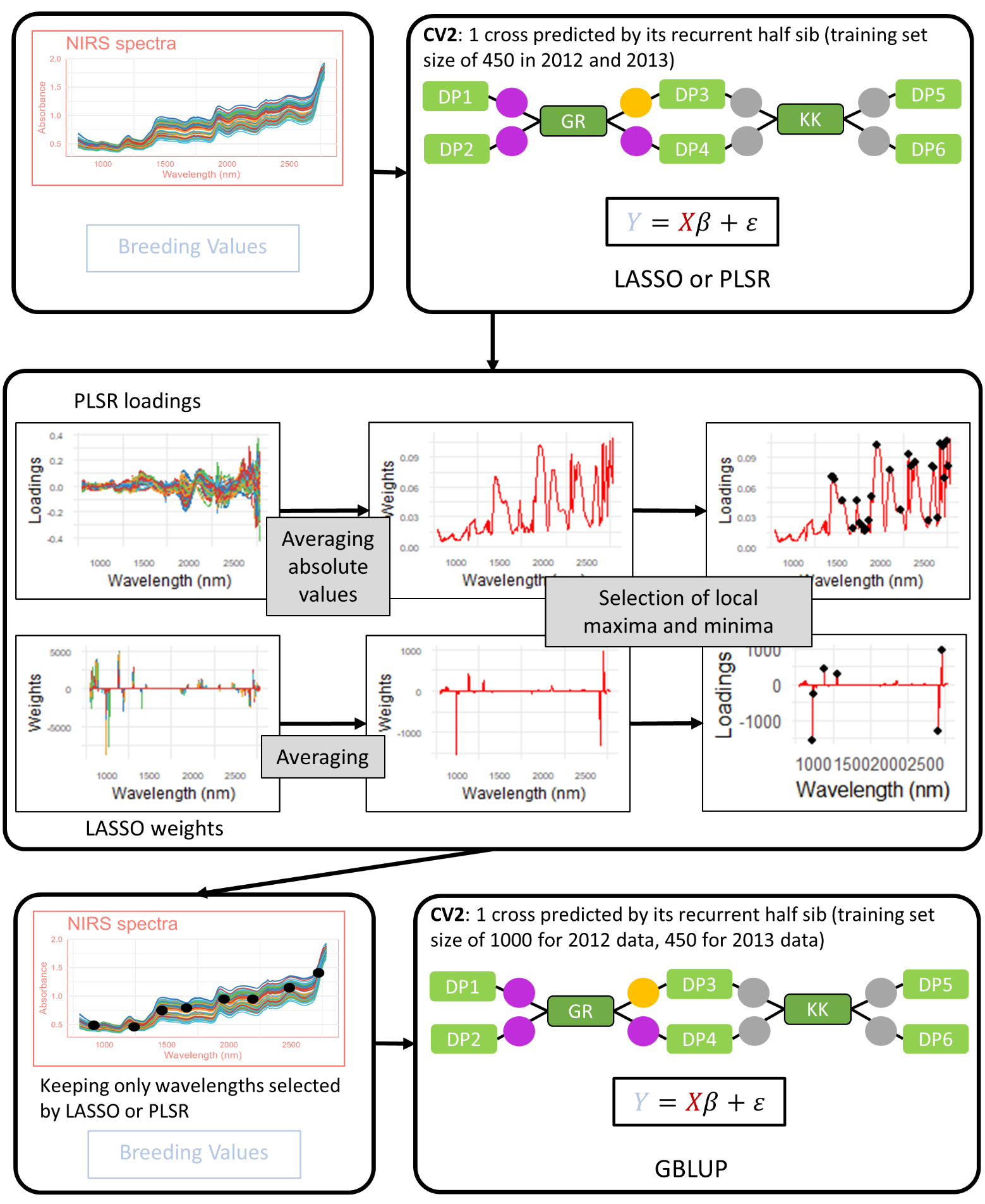

**Fig. S1: Schematic representation of the wavelength selection procedure**
Loadings and weights graphs are represented for PH (plant height) with LASSO and PLSR models trained on 2012 data in CV2 with a training set size of 450 with spectra pre-processed with SNV.

|  | 2012 | | | | 2013 | | | |
| --- | --- | --- | --- | --- | --- | --- | --- | --- |
|  | CZ1 | CZ2 | SB1 | SB2 | CZ1 | CZ2 | SB1 | SB2 |
| PAN | 0.17 | 0.18 | 0.23 | 0.21 | 0.19 | 0.19 | 0.19 | 0.21 |
| PED | 0.27 | 0.34 | 0.30 | 0.27 | 0.35 | 0.33 | 0.27 | 0.23 |
| STEM | 0.36 | 0.35 | 0.42 | 0.33 | 0.29 | 0.27 | 0.34 | 0.33 |
| PH | 0.28 | 0.26 | 0.30 | 0.23 | 0.24 | 0.22 | 0.27 | 0.23 |
| YIELD | 0.40 | 0.43 | 0.70 | 0.39 | 0.50 | 0.55 | 0.50 | 0.57 |
| FLAG | 0.08 | 0.09 | 0.09 | 0.08 | 0.08 | 0.09 | 0.10 | 0.08 |
| NIN | 0.16 | 0.17 | 0.20 | 0.24 | 0.15 | 0.16 | 0.17 | 0.20 |

**Table S1: Coefficient of variation of phenotypic measurement for each trait in each environment**

For each trait and each environment, coefficient of variation was computed as the standard error divided by the mean.

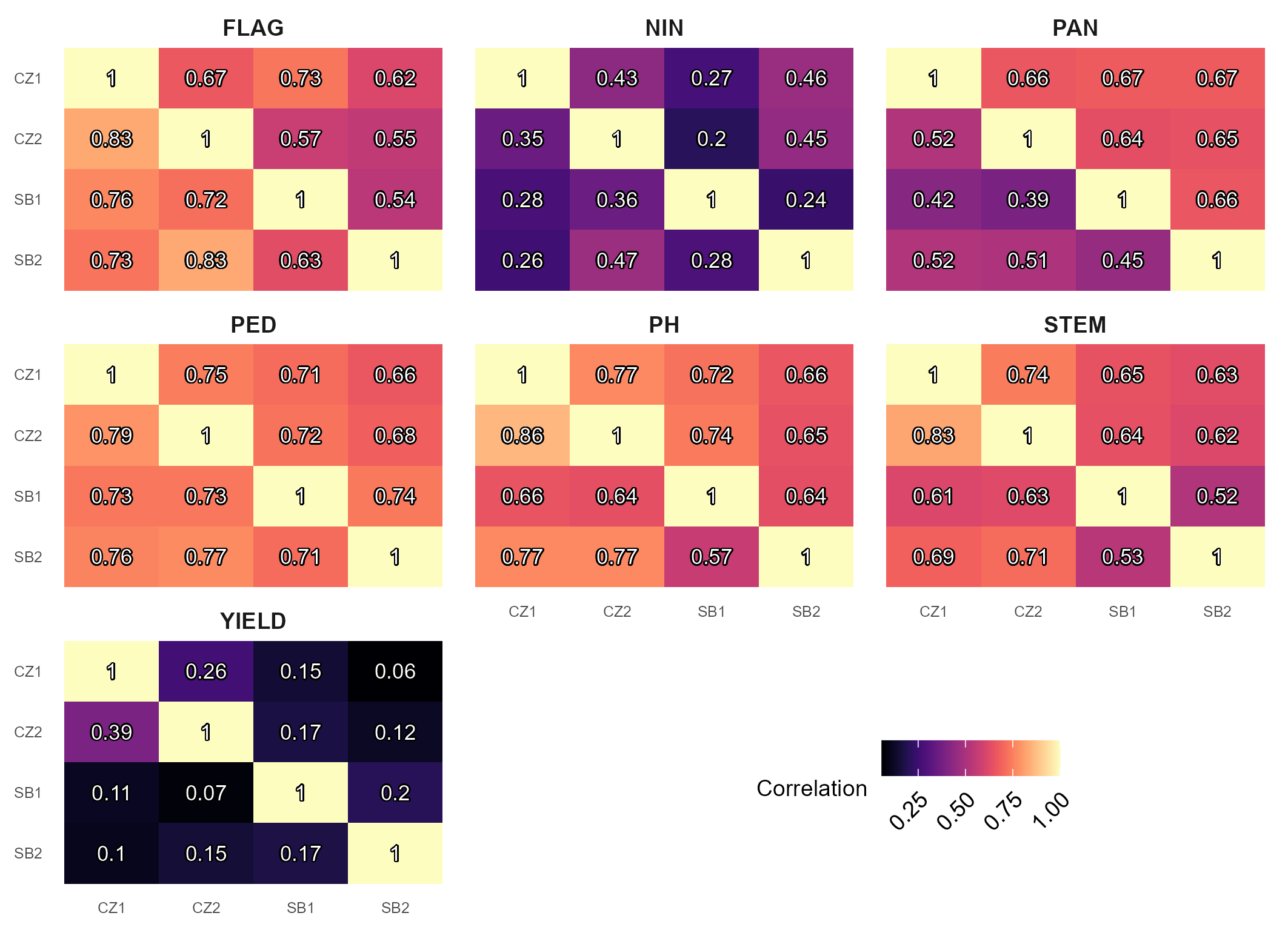

**Fig. S2: Correlation between environments for each trait in each year.**

Numbers are the Pearson correlation coefficient between values of different environments. Tiles under the diagonal are correlations for 2012, and tiles above the diagonal are correlation for 2013.

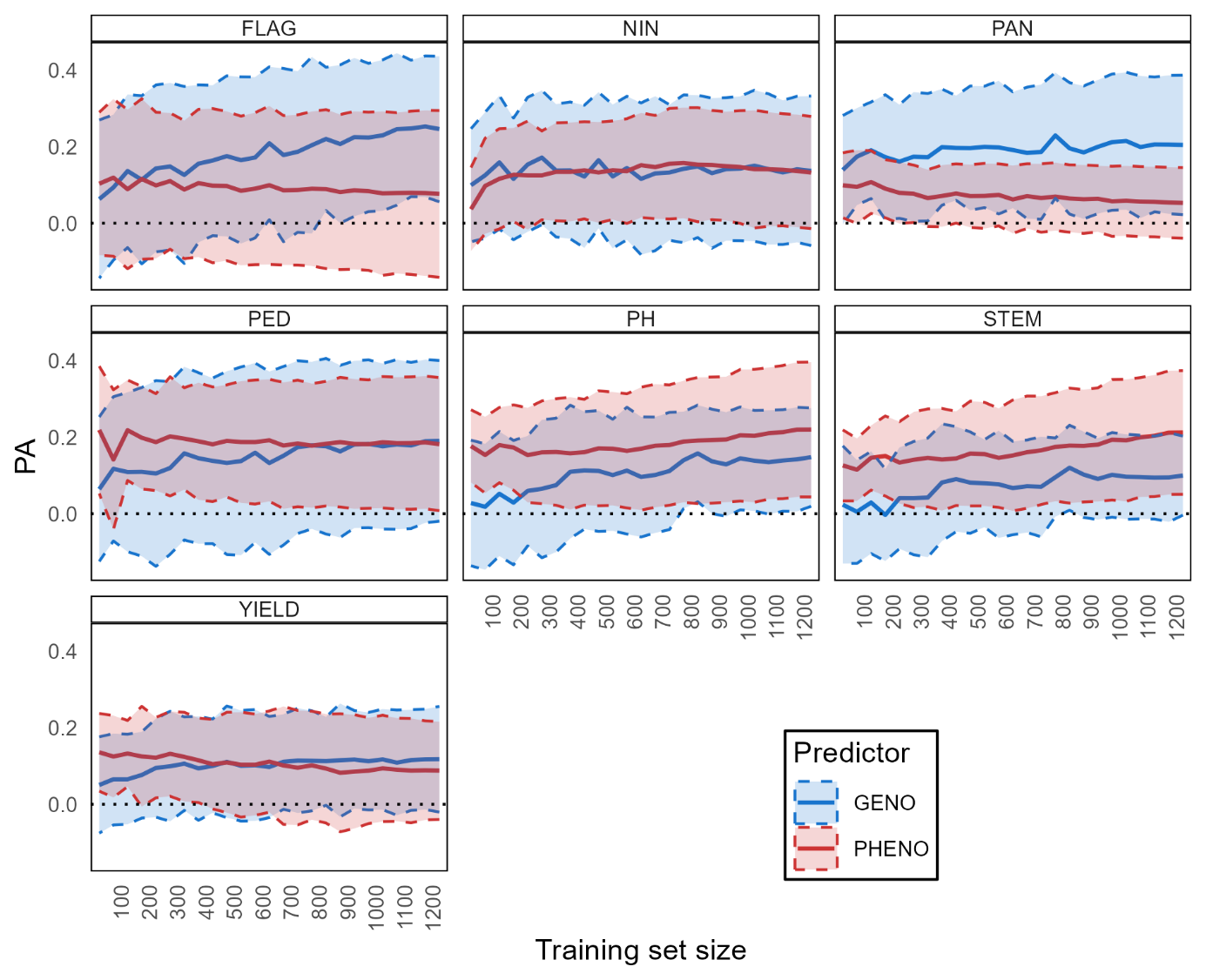

***Fig. S3: Effect of training set size on predictive abilities***
Predictive abilities were calculated with 2012 data, using GBLUP in the CV3 scenario and spectra pre-processed with SNV. Horizontal dotted line is at the 0 value. FLAG = flag leaf appearance, NIN = number of inter-nodes, PAN = panicle length, PED = peduncle length, PH = plant height, STEM = stem length, YIELD = yield. Full lines represent the mean PAs and dotted lines represent the standard error around the mean PA.

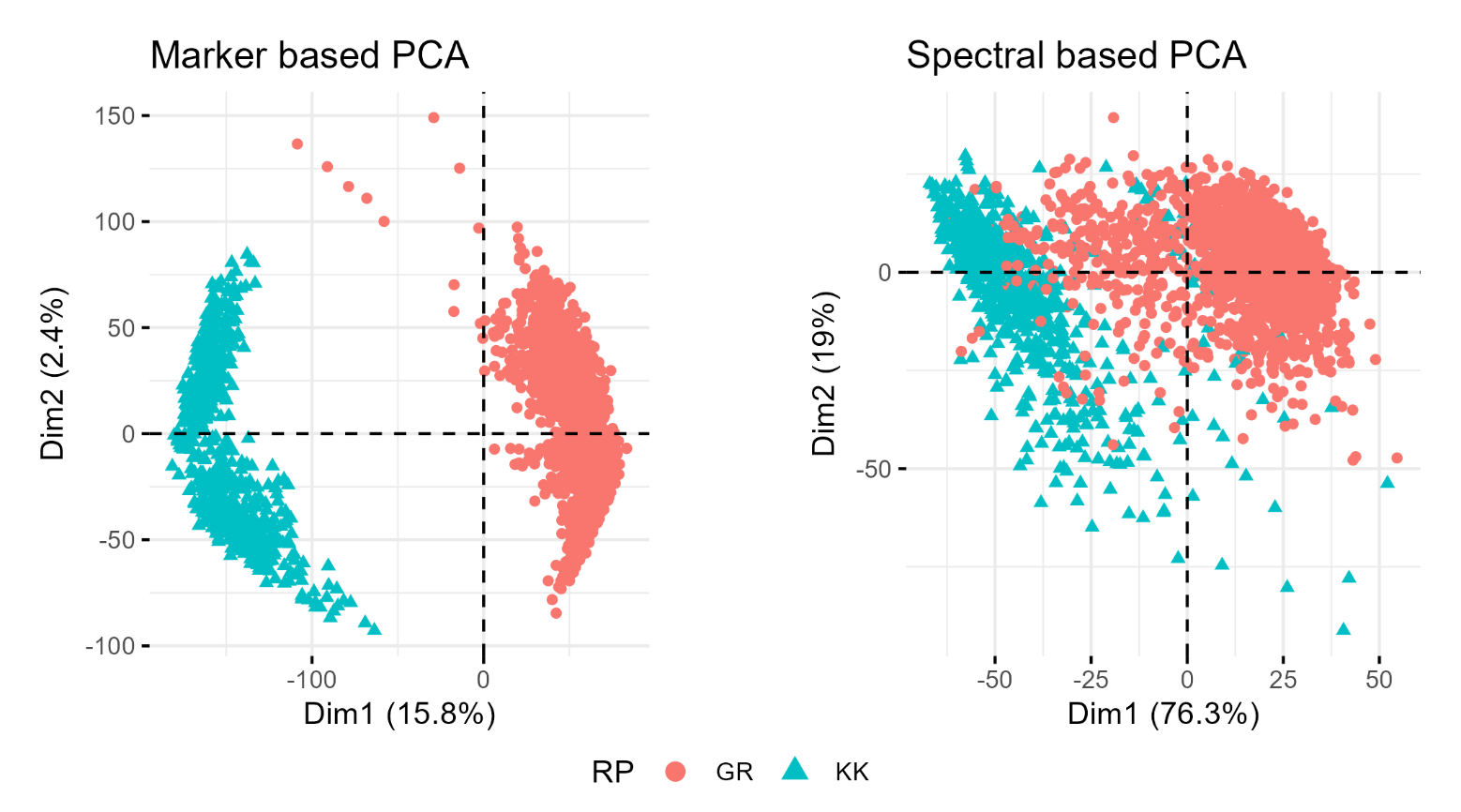

**Fig. S4: PCA of BCNAM based on marker or spectral data**

The two first components of principal component analysis are represented. PCA on genotype data was carried out on every SNP available, and PCA on SNV pre-processed spectra was carried out on all wavelengths available. RP = recurrent parent.

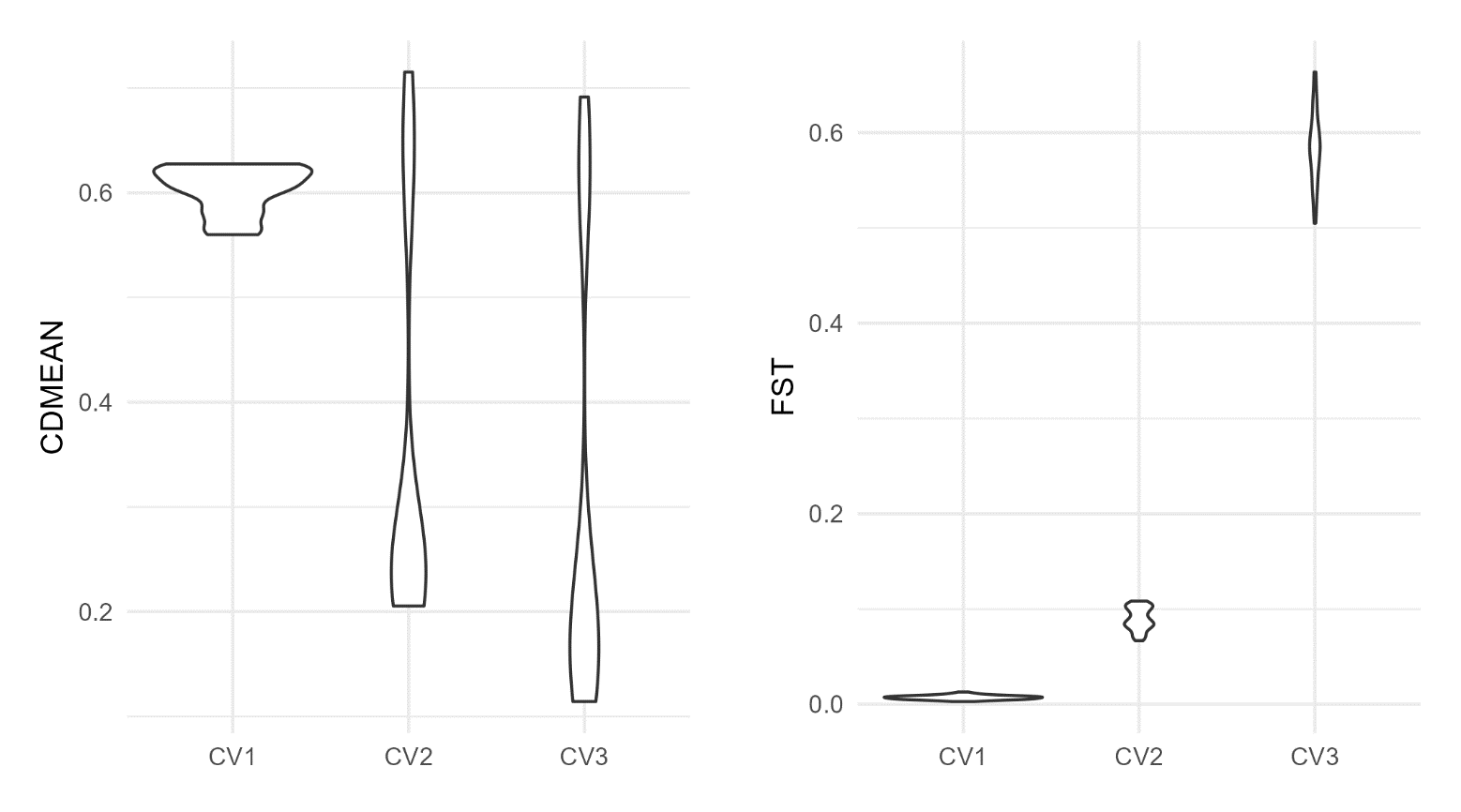

**Fig. S5: relatedness between CVs**

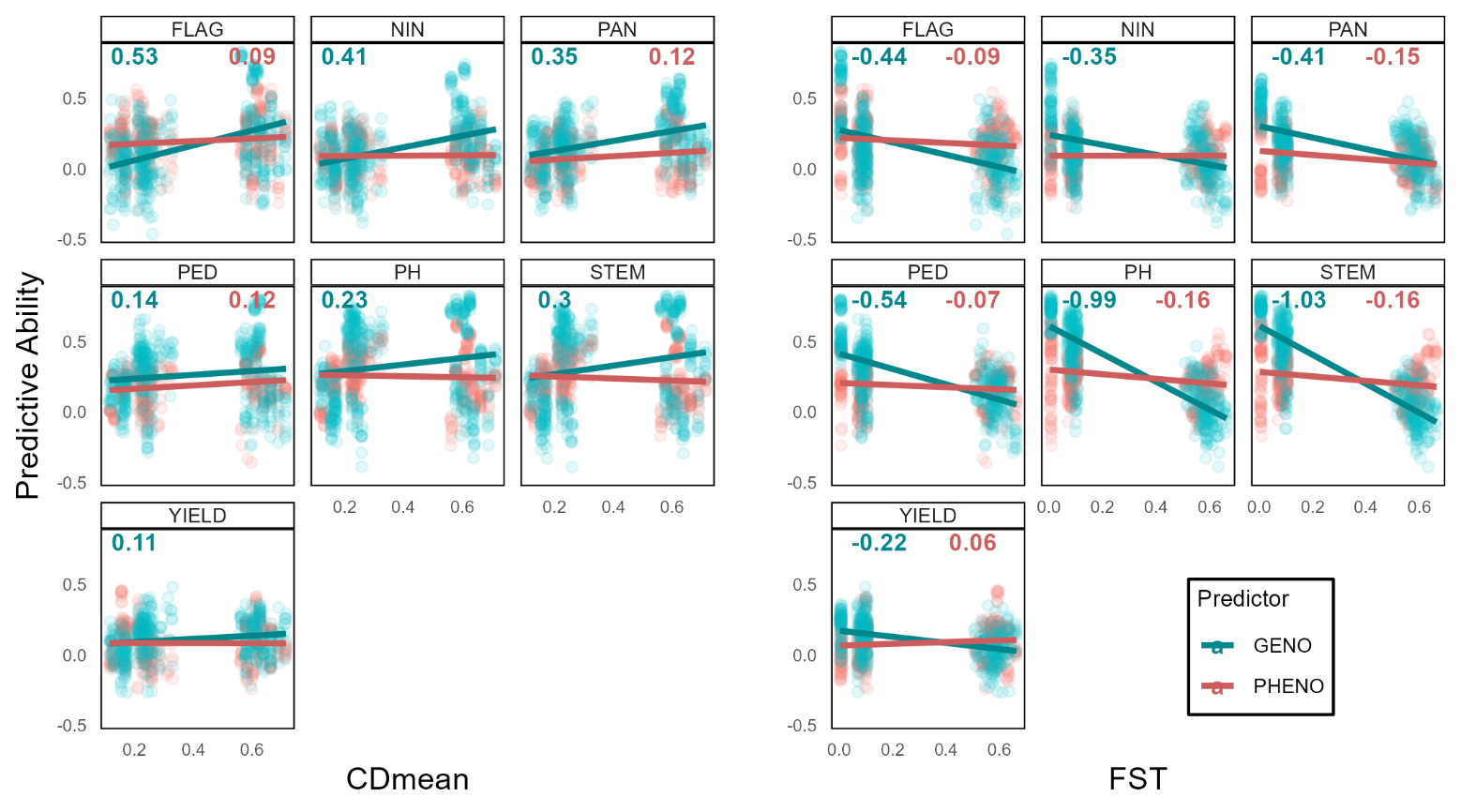

**Fig. S6: Effect of CDmean and Fst between training and validation sets on predictive abilities**

CDmeans and Fst were calculated on the same partitions used in CV1, CV2, and CV3. Numbers are the slopes of the linear regressions, and were displayed only when significantly different from 0 with a threshold of 5%.

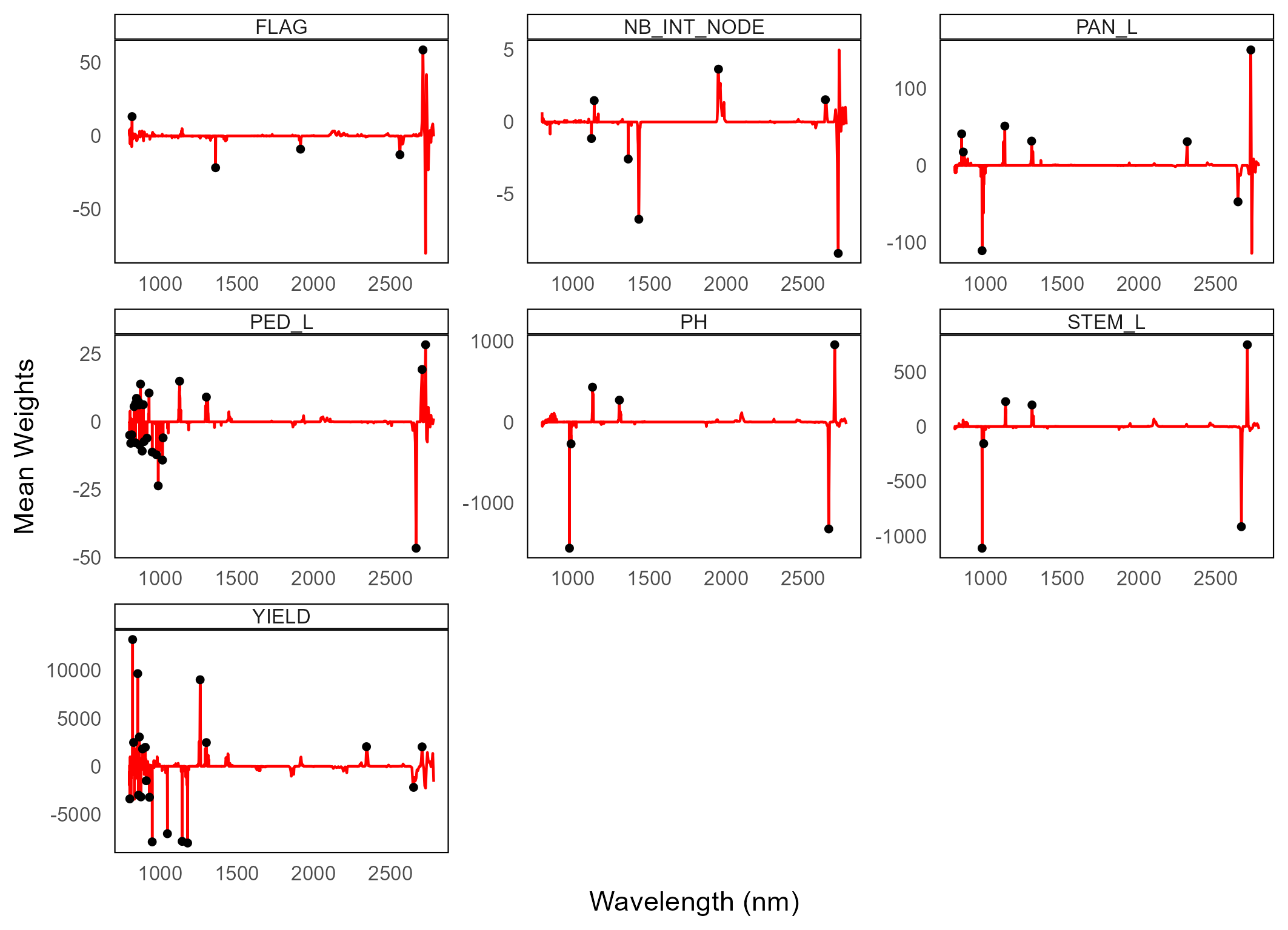

**Fig. S7: wavelengths selected by LASSO on 2012 data**
The red line is the mean of weights retrieved from LASSO models trained on 2012 data in CV2 with a training set size of 450 with spectra pre-processed with SNV. Black dots mark the wavelengths that were selected.

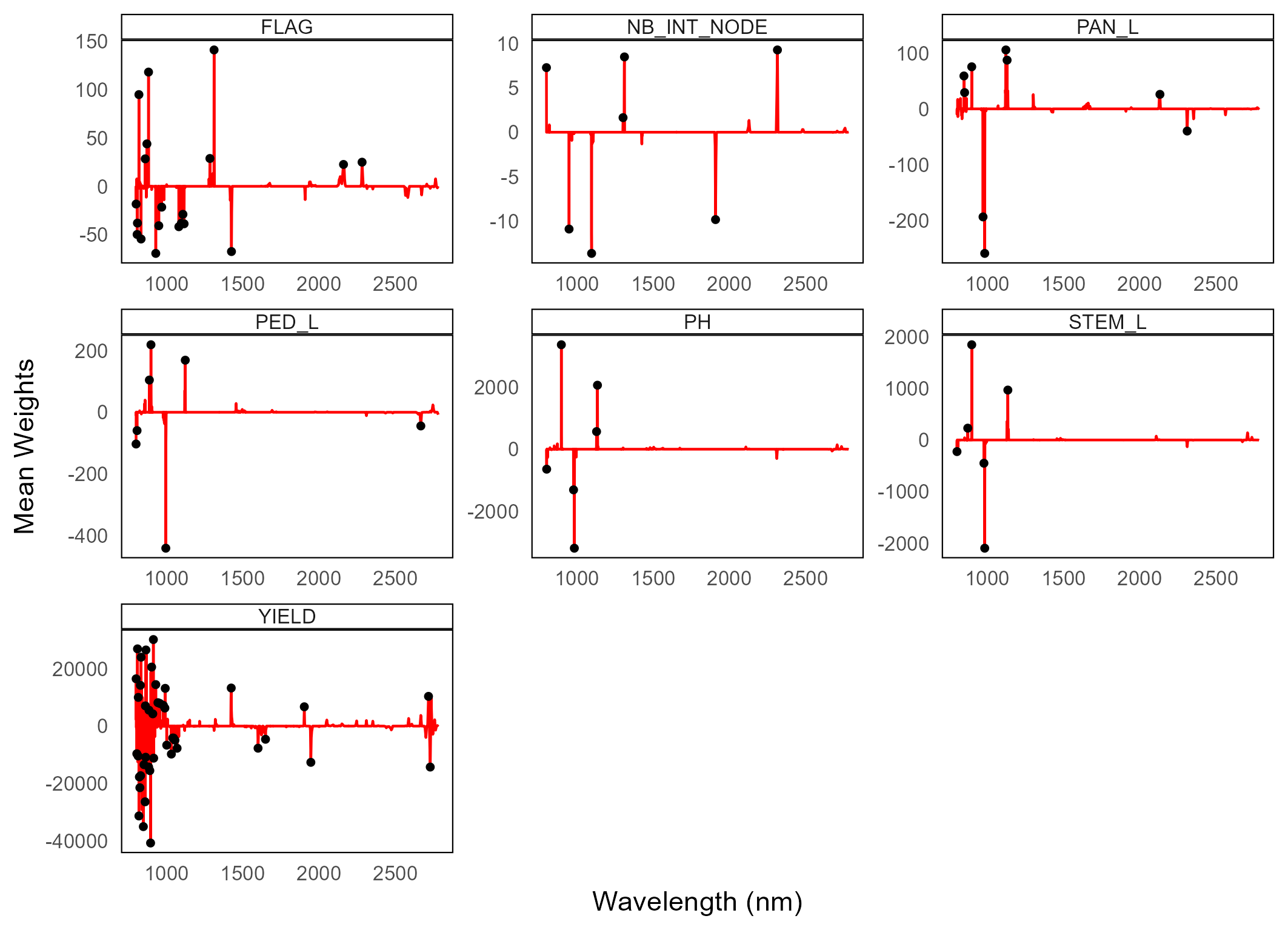

**Fig. S8: wavelengths selected by LASSO on 2013 data**
The red line is the mean of weights retrieved from LASSO models trained on 2013 data in CV2 with a training set size of 450 with spectra pre-processed with SNV. Black dots mark the wavelengths that were selected.

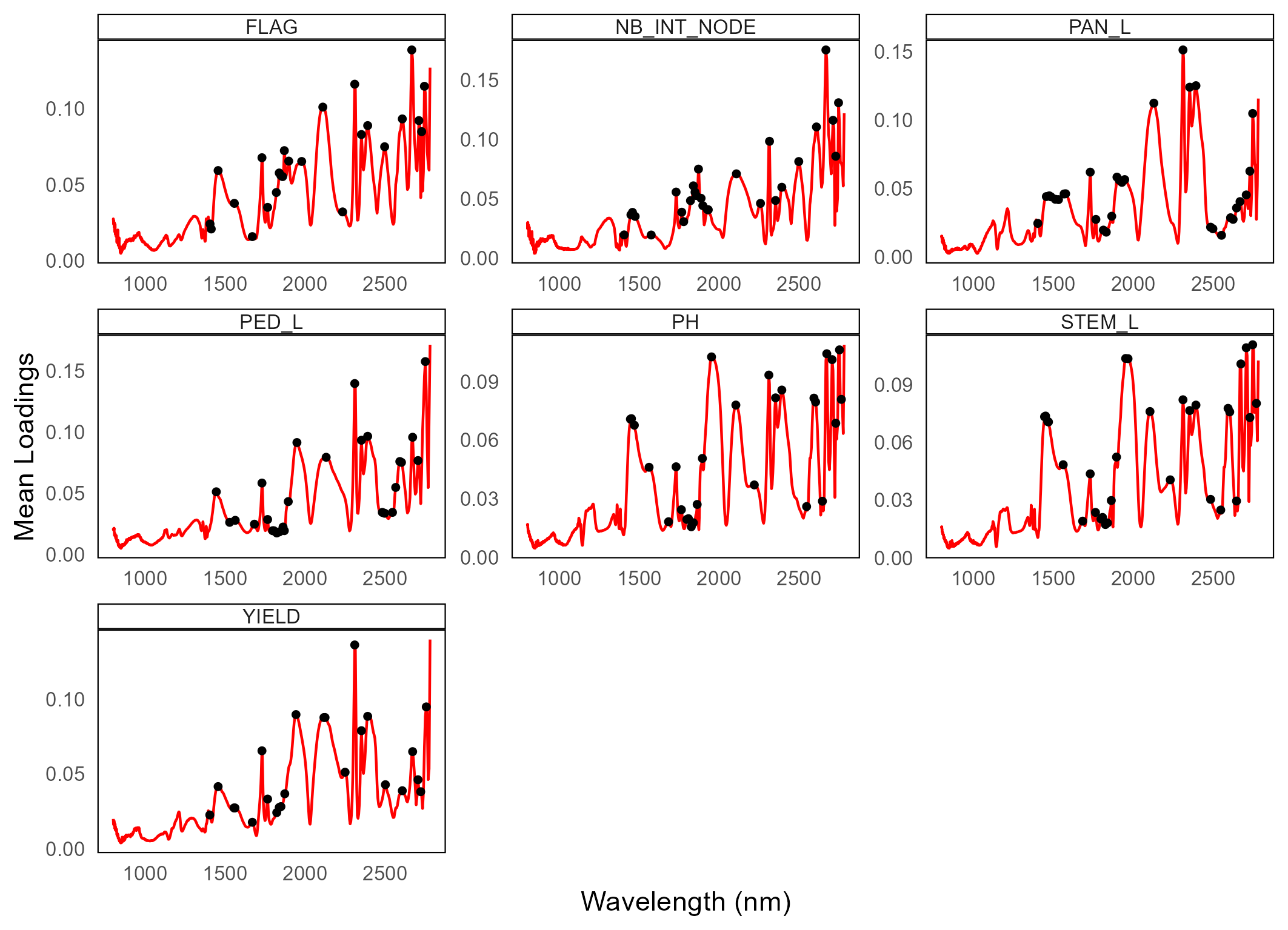

**Fig. S9: wavelengths selected by PLSR on 2012 data**
The red line is the mean of weights retrieved from PLSR models trained on 2012 data in CV2 with a training set size of 450 with spectra pre-processed with SNV. Black dots mark the wavelengths that were selected.

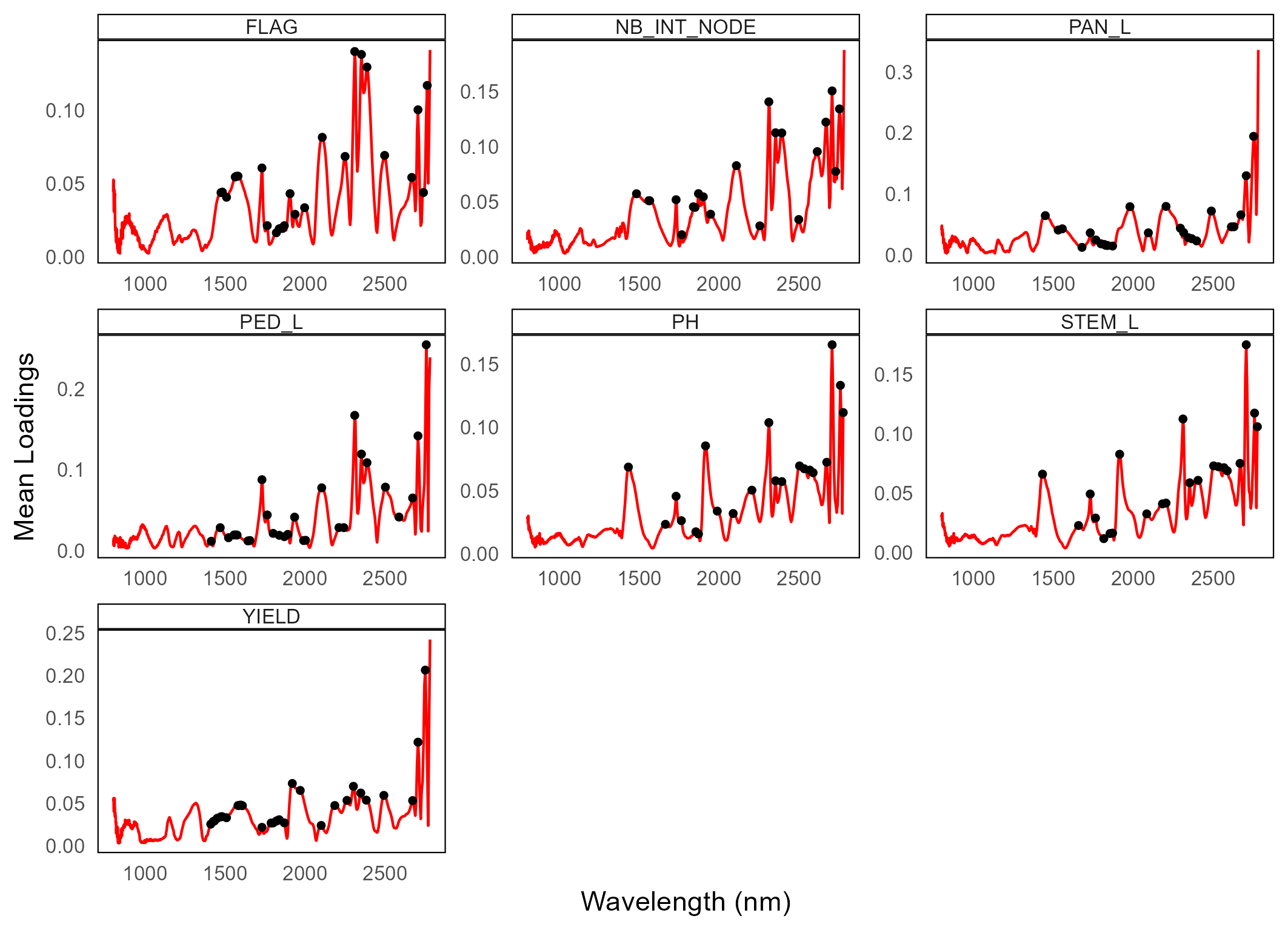

**Fig. S10: wavelengths selected by PLSR on 2013 data**
The red line is the mean of weights retrieved from PLSR models trained on 2013 data in CV2 with a training set size of 450 with spectra pre-processed with SNV. Black dots mark the wavelengths that were selected.
